# A new Petri Net Model of Working Memory to Predict the Root Causes of Attention Deficit Disorder Symptoms

**DOI:** 10.1101/2020.12.10.419093

**Authors:** Golnaz Baghdadi, Ali Doustmohammadi, Ainaz Jamshidi, Farzad Towhidkhah

## Abstract

Working memory is a system that helps us to store, retrieve, and manipulate information for a short period. The improper function of working memory is highly reported in people with attention deficit disorder. Attention deficit disorder is one of the most common disruptive behavioral disorders in children. Finding the actual reasons that may lead to inattentive symptoms is still an enigma for scientists. In this study, a model was proposed to show the flow of information through sensory, long-term, and working memory based on the Petri net approach. A new “selective updating” mechanism is also suggested. It is speculated that the central executive part of working memory uses this mechanism for updating the less important content with new incoming essential inputs. The analysis of the proposed model shows how an abnormality in the time setting or unexpected delays in information transmission may lead to some symptoms of inattention. These predictions about the possible causes of inattentive symptoms would be valuable for psychologists to find new possible treatments. This study also illustrates the great potential of Petri net approach for modeling and analysis of biological systems.

## 1. Introduction

Attention deficit disorder (ADD) is one of the most common neurobehavioral disorders developed in children, and its symptoms often last into adulthood (Biederman, 1998; Lofthouse, Arnold, Hersch, Hurt, & DeBeus, 2011). Finding the causes of these symptoms is one of the open problems in studies. Memory and attention are two terms that are seen together in the literature (Awh, Vogel, & Oh, 2006; Chun & Turk-Browne, 2007; Kuhl & Chun, 2014). In that, memory impairment may lead to attention deficit symptoms, and attention deficit disorder may prevent proper memory formation. Memory is conceptually divided into three stages of sensory, short-term, and long-term. Sensory memory (SM) holds sensory inputs (such as visual, auditory, etc.) temporarily (for less than one second) (Sperling, 1960). Short-term memory (STM) is a limited capacity store (about 7 + 2 units) (Miller, 1956). Long-term memory (LTM) is a more permanent, approximately limitless store, containing all our knowledge and memories (Cowan, 2008). All the mentioned types of memory are passive. That is, they are a place to store the data, and they do not entail the manipulation or organization of their information. Miller et al. (1960) used the term “working memory” instead of STM (Baddeley, 2003). Working memory (WM) is a theoretical framework that refers to structures and processes that allow us to hold and manipulate the information online. Therefore, while WM contains STM components, the concept of STM is distinct from WM. Working memory is necessary for doing complex tasks such as learning (Peijnenborgh, Hurks, Aldenkamp, Vles, & Hendriksen, 2016), reading comprehension (Nouwens, Groen, & Verhoeven, 2017), mathematical mental operations (Klados, Simos, Micheloyannis, Margulies, & Bamidis, 2015), and reasoning (Aries, Groot, & van den Brink, 2015) in the presence of distractors or interfering operations.

A strong relationship between WM capacity and the ability to attention control has been reported so far (Awh et al., 2006; Fougnie, 2008). Low WM capacity is one of the characteristics of people with ADD (Bayerl et al., 2010; Borella, Carretti, Riboldi, & De Beni, 2010; Dowson et al., 2004; Engle, Kane, & Tuholski, 1999; Fassbender et al., 2011; Klingberg et al., 2005; Klingberg, Forssberg, & Westerberg, 2002; Kobel et al., 2010; Kobel et al., 2009; Martinussen, Hayden, Hogg-Johnson, & Tannock, 2005; Redick & Engle, 2006; Schweitzer et al., 2004). However, WM is not a passive memory that only needs enough capacity to do its function properly. Working memory includes some mechanisms to import, remove, and update information selectively (Oberauer, Souza, Druey, & Gade, 2013). Therefore, it can be valuable to investigate the other possible problems of WM that may lead to ADD symptoms. Thus, we were motivated to provide a computational model of WM and its related components. Computational models can combine physiological and psychological knowledge with computational theories. This combination can provide a tool to investigate the system under some specified conditions and can predict the possible behaviors of the system in these circumstances. So far, various computational models have been proposed for working memory. These models have described WM from different aspects, such as the mechanism of encoding the information (Henson, 1998; Oberauer, 2009; Oberauer & Lewandowsky, 2011; Oberauer et al., 2013), role of attention (Baghdadi, Towhidkhah, & Rostami, 2017; Oberauer, 2009; Oberauer & Lewandowsky, 2011; Oberauer et al., 2013), rehearsal loop (Burgess & Hitch, 1992; Page & Norris, 1998), dealing with interferences (Lewandowsky & Farrell, 2005), learning and retrieving information (Brown, Hulme, & Preece, 2000), role of the involved cortical and subcortical regions (Ashby, Ell, Valentin, & Casale, 2005; R. O’Reilly, 2003; R. C. O’Reilly, Braver, & Cohen, 1999), the capacity limits (Bays, 2018), and controlling the entrance of information (Frank, Loughry, & O’Reilly, 2001; R. C. O’Reilly et al., 1999; R. C. O’Reilly & Frank, 2006).

Several conceptual models and theories were also proposed to show the characteristics of WM from different aspects (for a review see (Oberauer et al., 2018)). We used some of them to form our proposed model that their details have been provided in the next sections. In this paper, we have tried to integrate and represent the mentioned WM concepts and theories using Petri net (PN) approach.

Petri net is a well-known computational modeling approach for discrete event systems (DES) such as manufacturing and industrial systems (Hrz & Zhou, 2007; Zhou & Kurapati, 1999) as well as biochemical processes (Blätke, Heiner, & Marwan, 2011). Discrete event systems are characterized by a set of states that can change with the occurrence of events at discrete times (Zhou & Kurapati, 1999). With this definition, most of the voluntary and involuntary mechanism of human behaviors in conditions such as professor-student interaction (Kouzehgar, Badamchizadeh, & Khanmohammadi, 2013), driving (Thiruvengada & Rothrock, 2007), and planning (Joo et al., 2013) can be categorized as DES. Memory is another biological system that events form its operational dynamic (Friedman & Johnson Jr., 2000; Missonnier et al., 2005). In the current research, we used PN approach to design a model of WM based on the available knowledge about its operation. The proposed model shows the connection between WM and its related components (i.e., SM and LTM), the flow of information through these components, and a possible WM selective updating method. This model provides some valuable predictions about probable causes of inattentive symptoms. These predictions are mainly focused on how changing the timing of different parts affects attention performance.

In the next section, PN approach is presented briefly. Then, observations of WM operation are described, followed by the PN representation in section 4. The analysis of the proposed model is reported in section 5. The paper is concluded in section 6 with some suggestions for future work.

## 2. Petri Net

Petri Net, since its origin by Carl Adam Petri in his 1962 Ph.D. dissertation on the study of the communication protocols between the components of a computer system, has been developed considerably (Silva, 2013). Petri Net has evolved into an elegant and powerful graphical modeling tool for asynchronous concurrent systems. Some advantages of PN are 1) its graphical nature enables easy visualization of complex systems; 2) a systematic and qualitative analysis is possible using well-developed techniques; 3) performance evaluations of systems are possible using timed PNs (Bouyer, Haddad, & Reynier, 2008; Damadi, Doustmohammadi, & Afshar, 1989; Murata, 1989).

The basic elements of a PN are places, transitions, tokens, and directed arcs to connect places with transitions. Tokens reside in places and move from one place to another when transitions are enabled in a system. Graphically, places, transitions, and tokens are respectively represented by ○, |, and •. The status of a place represents resources, the firing of a transition represents the occurrence of an event, and tokens represent the availability of resources in the system. The distribution of tokens in different places is called marking. The changes in marking can exhibit the dynamic of the system. In timed PN, solid and hollow rectangles are used for immediate and timed transitions, respectively. An immediate transition is enabled and fired as soon as all of its input places have sufficient tokens. Similarly, a timed transition is enabled as soon as all of its input places have enough tokens. However, its firing is delayed by the time attributed to the transition (Damadi et al., 1989). Transitions with inhibitory arcs are enabled only when the input places are empty. The above definitions have been summarized in Fig. 1.

**Fig. 1.**
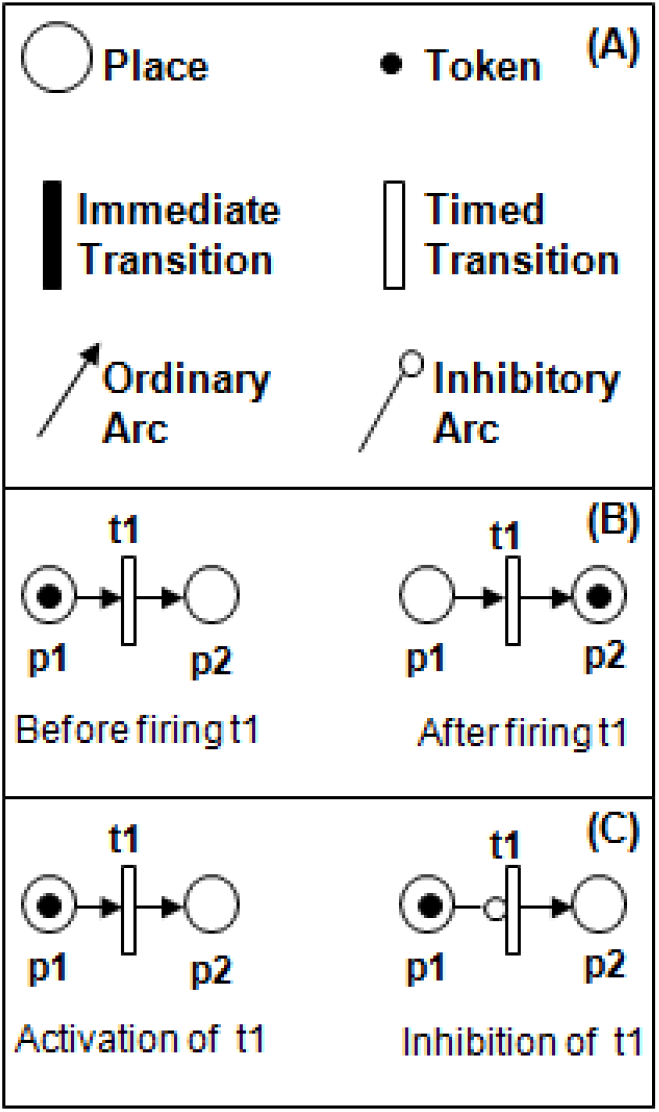
Graphical representation of the basic elements in PN

## 3. Working Memory Models

As mentioned in the introduction, SM, STM, WM, LTM are different types of memory that are distinguished based on their spatial and temporal limitations. Sensory memory can be considered as a buffer to hold incoming stimuli for about one second (Sperling, 1960). Long-term memory preserves information from a few days to decades. Short term memory keep information (about 7±2 units of information: the Miller’s magical number) received from SM for about 10-15 seconds (without any rehearsal) (Peterson & Peterson, 1959). During this time, WM actively protects the relevant information and tries to remove interfering disturbances. Therefore, STM and WM are not completely distinct from each other. Short-term memory is a limited capacity buffer to keep required information of SM and LTM, while WM actively protects and processes them. In 1968, Atkinson and Shiffrin proposed a conceptual model (Fig. 2) to show the relation between short term (working) memory, SM, and LTM (Atkinson & Shiffrin, 1968). In this model, “Input” refers to the environmental sounds, pictures, smells, etc. that are received by our sensors.

**Fig. 2.**
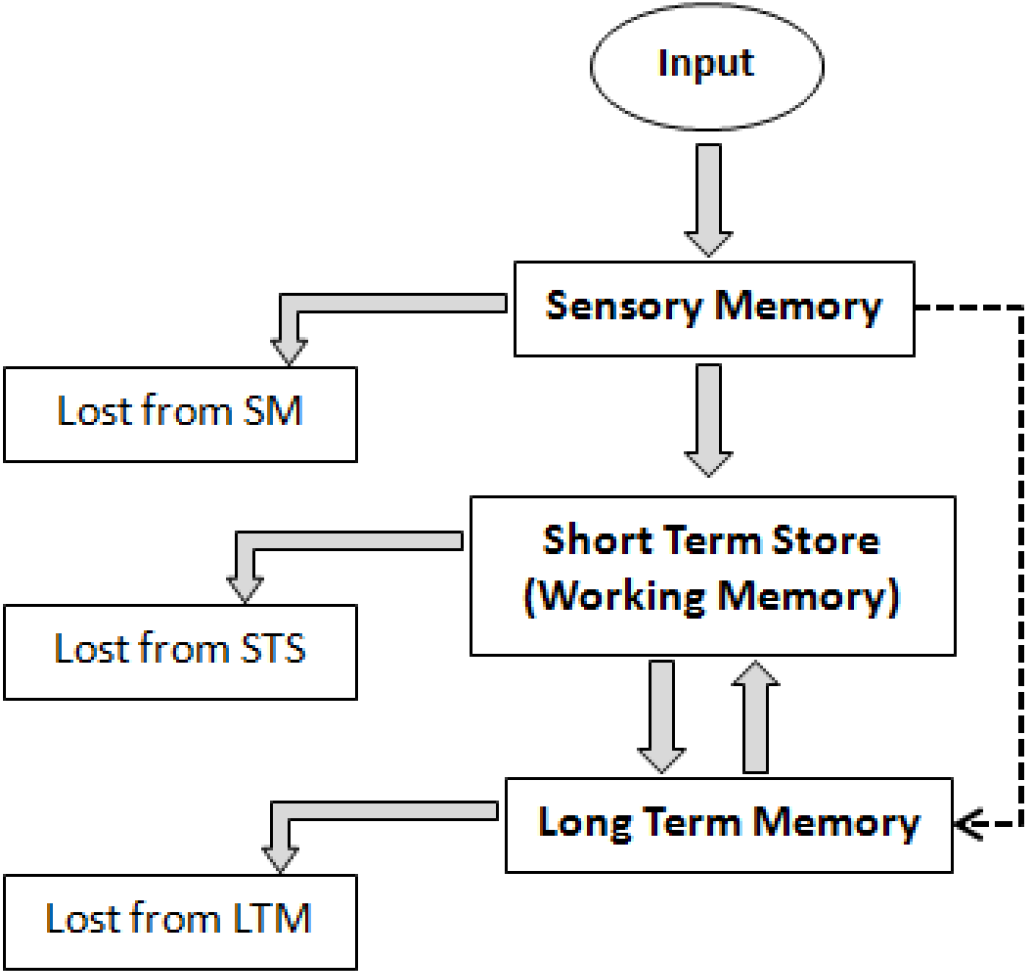
Short term (working) memory and its relation with sensory memory and long term memory; proposed by Atkinson and Shiffrin, 1968.

Although Miller et al. used the term “working memory” in the 1960s for the first time, it became much more dominant by the multi-component WM model of Baddeley and Hitch in 1974. This model is modified later by Baddeley in 2000 (Fig. 3) (Baddeley, 2000).

**Fig. 3.**
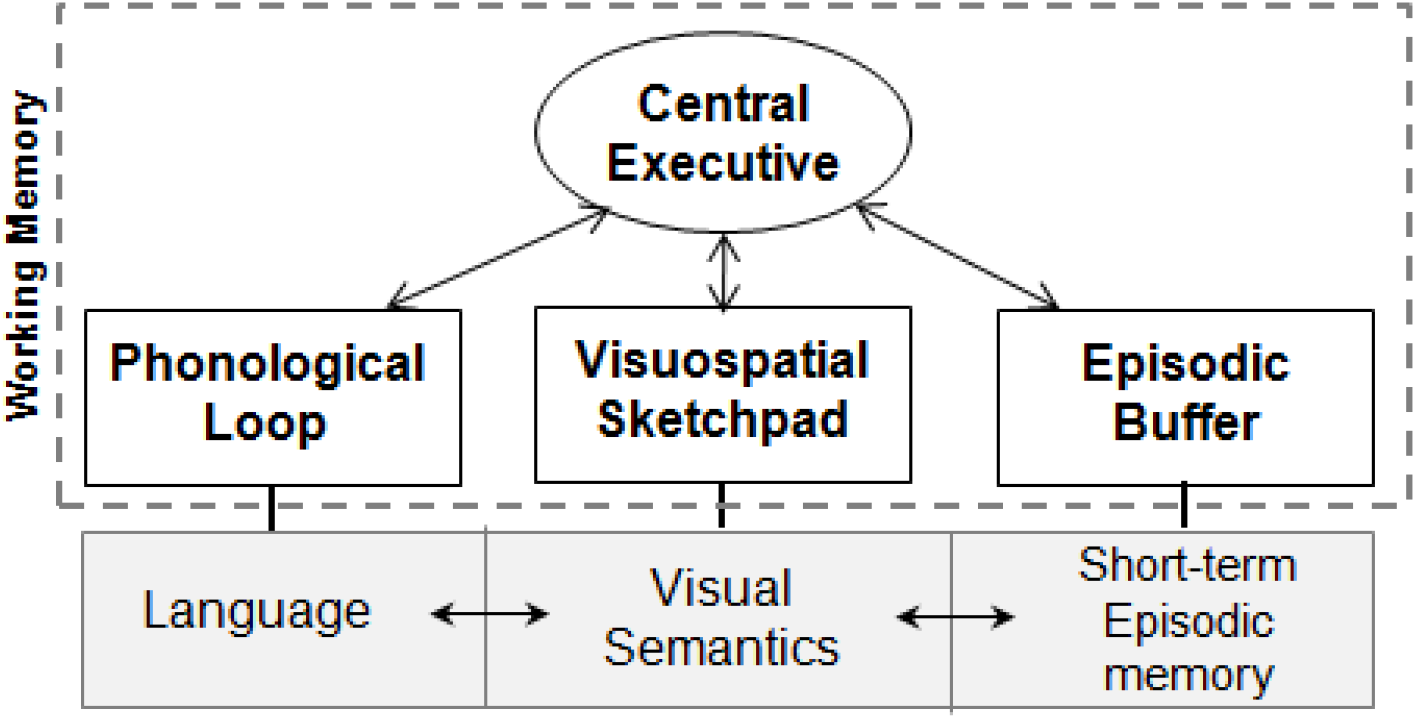
The revised model of working memory proposed by Baddeley in 2000

This revised version consists of a master subsystem “Central executive” that controls three slave parts: “Phonological loop,” “Visuospatial sketchpad,” and “Episodic buffer.” The intelligent part of this model is central executive, and other three slave subsystems hold different kinds of information with a relatively little processing. Phonological loop and visuospatial sketchpad subsystems are respectively considered for the temporary storage of speech and visual information. Episodic buffer stores a combination of phonological and visual information coming from SM or LTM (Baddeley, 2000). Baddeley divided the phonological loop into two subsystems: “Phonological store” and “Articulatory loop.” The first subsystem stores phonological information, and the second prevents the decay of information by continuous repetition of the content in a “rehearsal loop” (Jarrold, Baddeley, & Phillips, 1999) shown in Fig. 4.

**Fig. 4.**
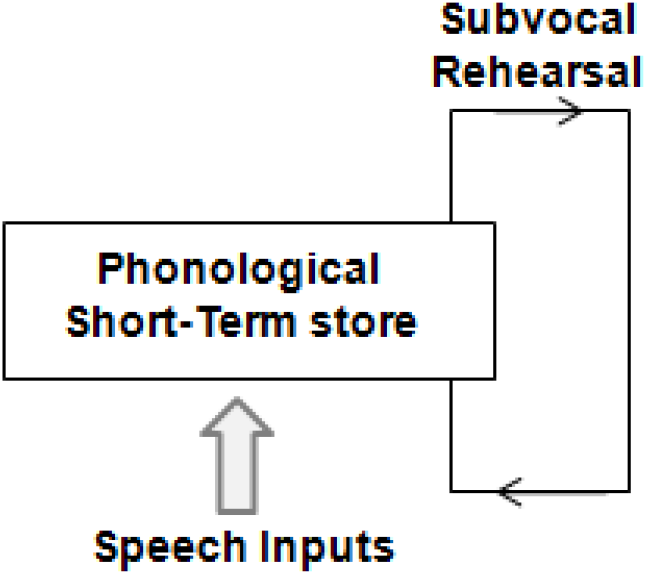
Baddeley’s model of the phonological loop (adapted from (Jarrold et al., 1999))

The central executive part controls the entrance of SM and LTM information into the short-term buffers. This part corresponds to the attentional control system in a WM model proposed by Oberauer (Oberauer, 2009; Oberauer et al., 2013). Central executive part or attentional control system inhibits the entrance of irrelevant information to the short-term storage (STS) buffers (Kofler, Rapport, Bolden, Sarver, & Raiker, 2009; Oberauer et al., 2013). The procedure of this inhibition is called “selective active gating mechanism” (Frank et al., 2001), which has been shown in Fig. 5. When the gate is open, sensory input is allowed to enter to the WM buffer. When the sensory input is irrelevant, the gate is closed to maintain the previously stored information. This mechanism guarantees the “robust maintenance” of relevant information in WM buffers (Frank et al., 2001).

**Fig. 5.**
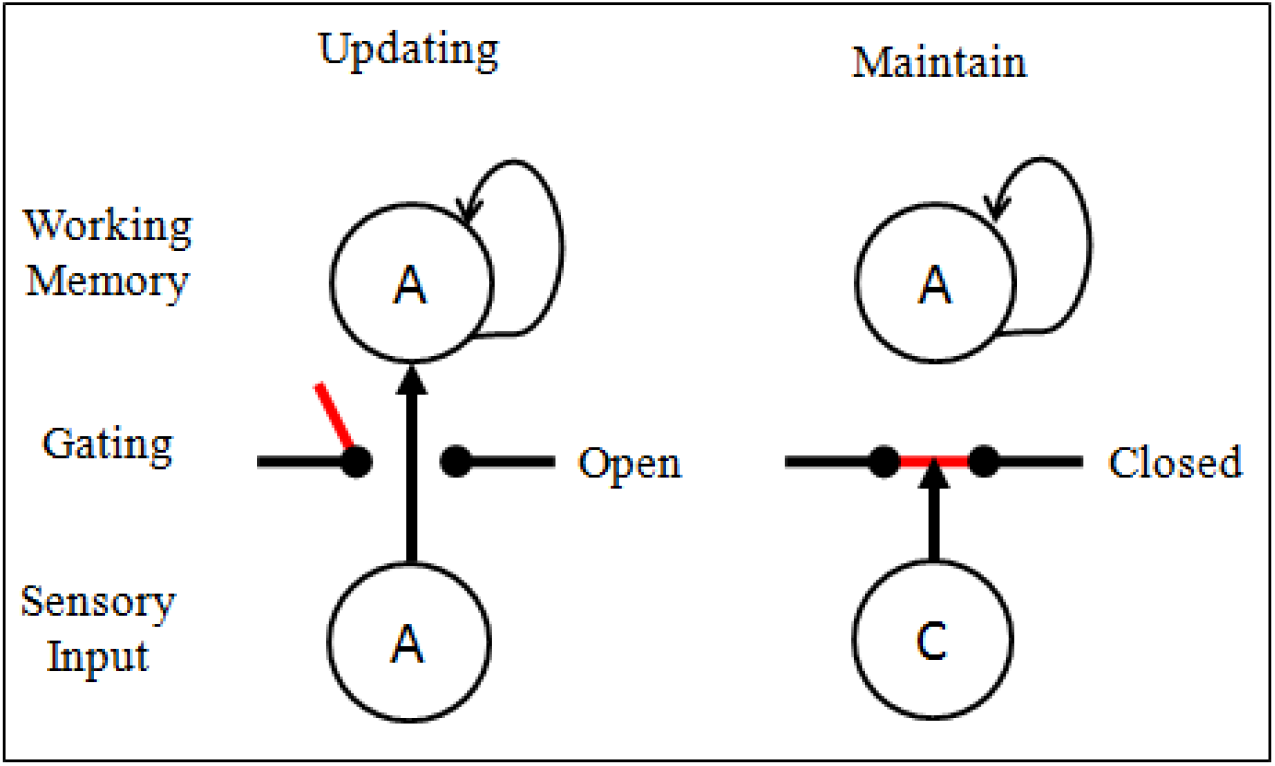
Active gating mechanism illustration (adapted from (Frank et al., 2001)): “A” is a relevant and “C” is an irrelevant input. Active gating mechanism inhibits the entrance of “C” to keep the relevant information “A.”

How the gate is opened or closed? Dopamine is a neurotransmitter that acts as a “gating signal” (Badre, 2012; Frank, 2005; Gruber, Dayan, Gutkin, & Solla, 2006; Jarrold et al., 1999; Kofler et al., 2009; R. C. O’Reilly & Frank, 2006; Redgrave & Gurney, 2006). Dopamine neurons encode the prediction error of rewarding outcomes (R. C. O’Reilly et al., 1999; Redgrave & Gurney, 2006). Some parts of the brain map dopamine to the “Go” and “No-Go” signals (Hulme, Roodenrys, Brown, & Mercer, 1995). Consequently, dopamine is released to allow the entrance of the information (the “Go” signal) when the input is diagnosed as relevant data. Otherwise, the irrelevant information is not allowed to enter to the WM buffer.

An interaction between the prefrontal cortex and basal ganglia is associated with the producing of the “Go/No-Go” signal (dopamine) (R. C. O’Reilly & Frank, 2006). A computational model was proposed to show this interaction (R. C. O’Reilly & Frank, 2006). This model works based on the reinforcement learning mechanism. In that, according to the obtained reward (internally or externally), the model learns to send “Go” or “No-Go” signals in expected times.

Considering the concepts and characteristics listed for WM through the mentioned conceptual and computational models, we proposed a new model for WM based on PN concepts. The proposed model, which represents an integration of the mentioned characteristics, is explained in the following sections.

## 4. The Proposed Model

The structural framework of our model is based on the Atkinson and Shiffrin (1968) model (shown in Fig. 2). We have considered all three slave subsystems of Baddeley’s model as a single short term storage buffer with limited capacity and a central executive part to control and manage the flow of inputs from and to this storage buffer. The control is done using a combination of the “active gating mechanism” (Frank et al., 2001), and our suggested selective updating procedure. Fig. 6 shows the block diagram of the proposed model.

**Fig. 6.**
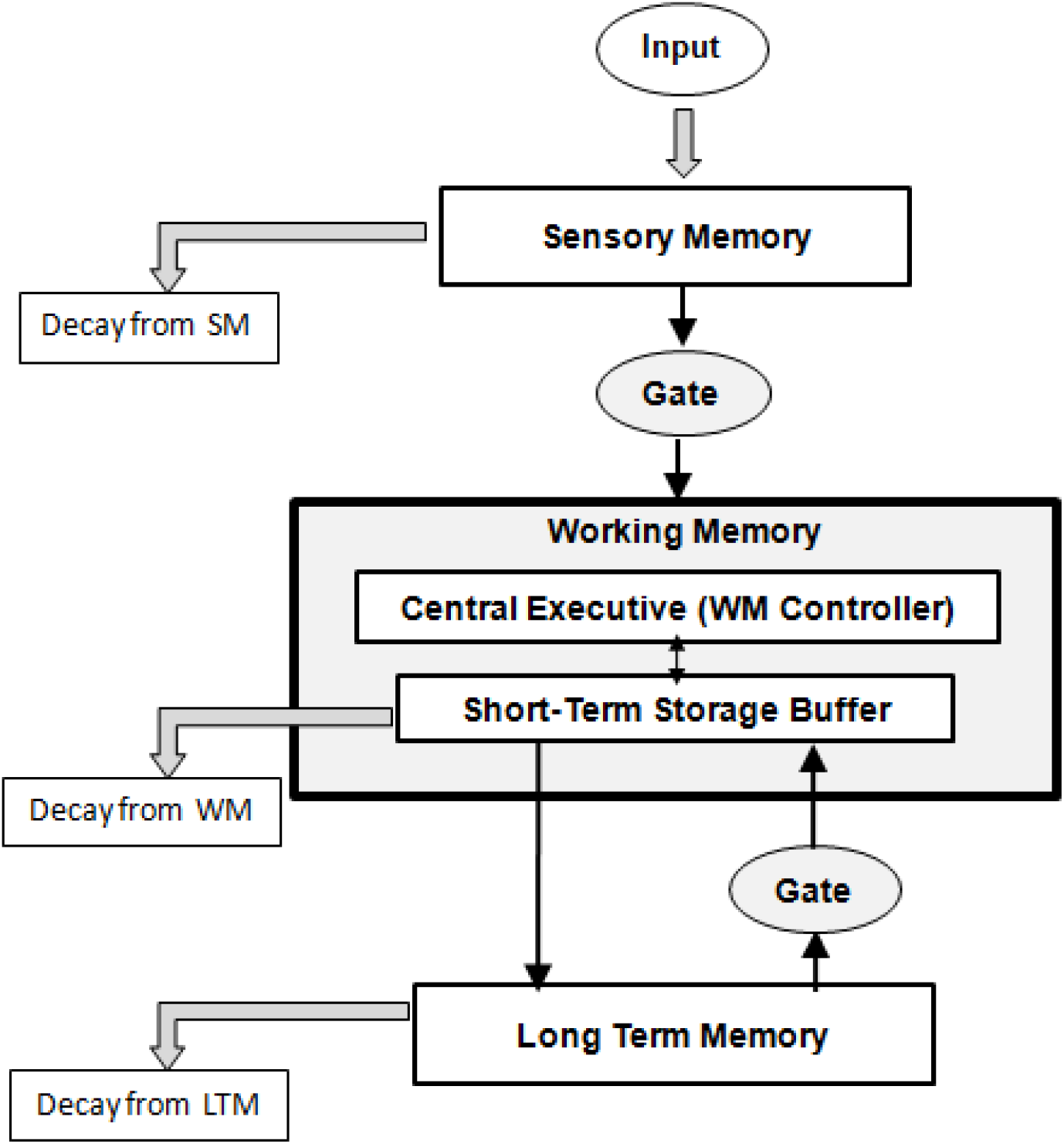
The block diagram of the proposed model

As shown in Fig. 6, short-term storage buffer of WM can receive input from SM and LTM. An active gate controls the inputs from these two memories (SM and LTM). Controlling the gates and contents of short-term storage buffer are performed in the central executive part. The Petri net presentation of this model has been illustrated in the next section.

### 3.1. Petri Net Representation of the Proposed Model

In this part, the PN graphical representation of our model is described, and its analysis is left for the next part. We have implemented our model by HPSim V1.1 software with the permission of its author Henryk Anschuetz. As mentioned before, the proposed model is a combination of the memory model of Atkinson and Shiffrin (Atkinson & Shiffrin, 1968), the WM model of Baddeley (Baddeley, 2000), and the proposed selective gating mechanism of Frank et al. (Frank et al., 2001). Considering the block diagram of the proposed model (Fig. 6), the PN representation of each part has been described in the following subsections.

### 3.2. Sensory Memory

According to Fig. 6, the first subsystem of the proposed model is SM that has been represented by the PN approach in Fig. 7.

**Fig. 7.**
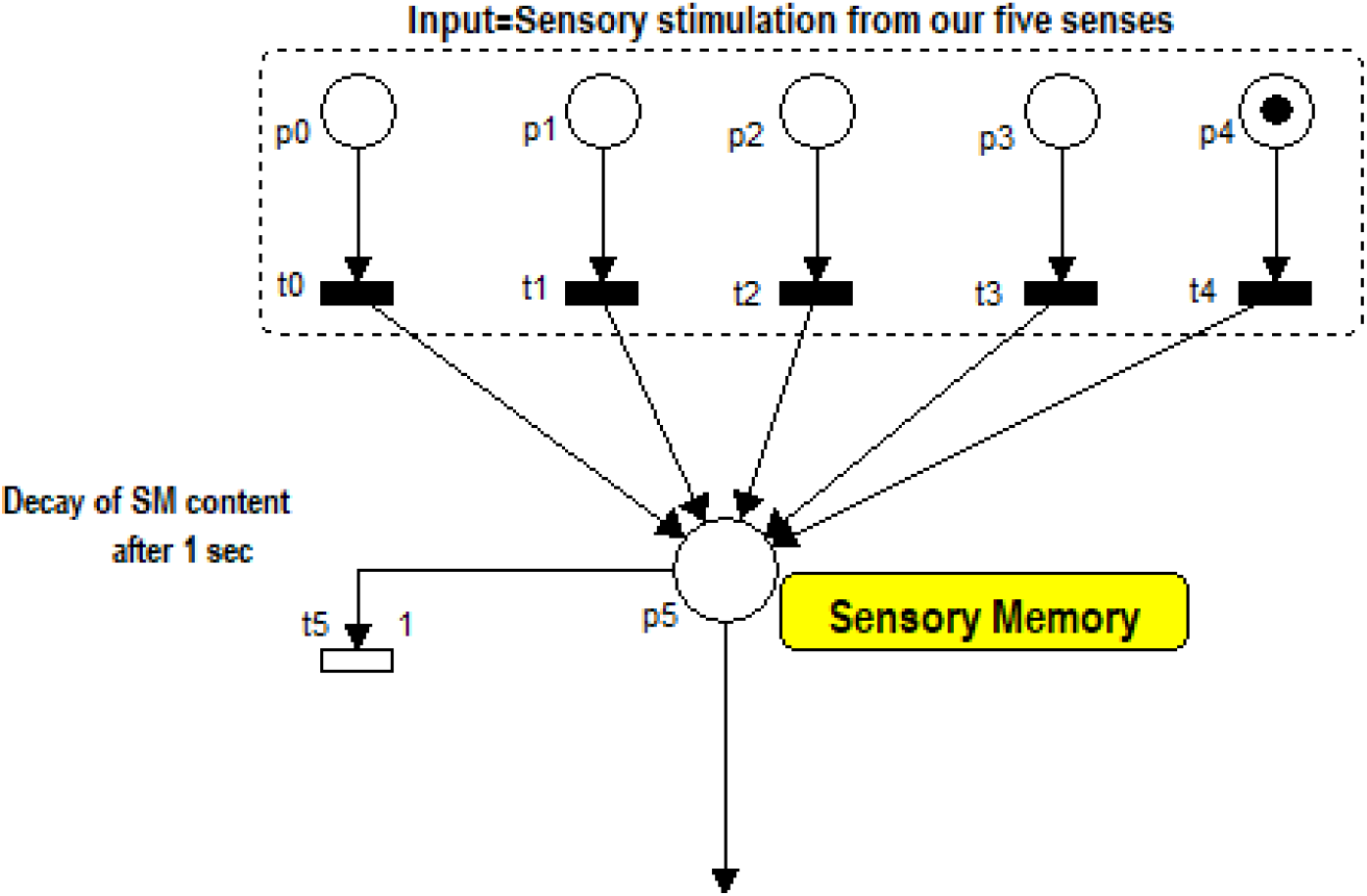
The Petri net representation of sensory memory in the proposed model. The tokens in p0-p4 are sensory inputs that respectively pass through immediate transitions t0-t4. Sensory memory, which has been represented by place p5, receives the incoming stimuli and waits for permission to send them to WM. This place can hold its contents for a short period (one second). After passing this time, they will be decay through timed transition t5 if the contents cannot find the chance to flow into WM.

As shown in Fig. 7, SM (place p5) holds the sensory stimuli that are received from our five senses: sight, hearing, smell, touch, and taste. Comparing Fig. 6 and Fig. 7, places p0-p4 represent the “input” and p5 represents the “sensory memory.” Tokens in places p0-p4 show the incoming stimuli. In this model, we have supposed that there is no time delay between stimulation incoming and its transition to SM. Therefore, this transition has been modeled by immediate transitions (t0-t4). However, in some disorders, there is an unexpected delay in the mentioned transitions that are not considered in the current study. In Fig. 7, the firing of t5 leads to the decay of SM content. Sperling in 1960 reported that SM could hold information for less than one second (Sperling, 1960). Therefore, the average firing time of t5 has been set on one second. If the SM content is transferred from SM to WM in less than one second, then t5 cannot fire since no token is available in p5. The flow of the relevant inputs from SM to WM is the next issue that has been described in the next subsection.

### 3.3. Active Gating

Central executive part has the most important role in controlling attention and flow of inputs to WM (Arias-Carrión & Pöppel, 2007; Baddeley, 2000). According to the previous discussion, “active gating” is one of the proposed mechanisms that helps the central executive part of WM to control the flow of input. The gate between SM and WM should be opened to allow the SM content to enter the STS buffer of WM and should be closed to avoid the entrance of distracting inputs. Two gates are considered in the proposed model: one of them controls the SM inputs to avoid the entrance of distracting stimuli (external noise), and the other located between LTM and WM to avoid the entrance of irrelevant data from LTM (internal noise). Modeling such a gate (shown in Fig. 5) can be performed by a transition that its activation depends on the existence of a token in a place. Fig. 8 shows the PN presentation of these two gates.

**Fig. 8.**
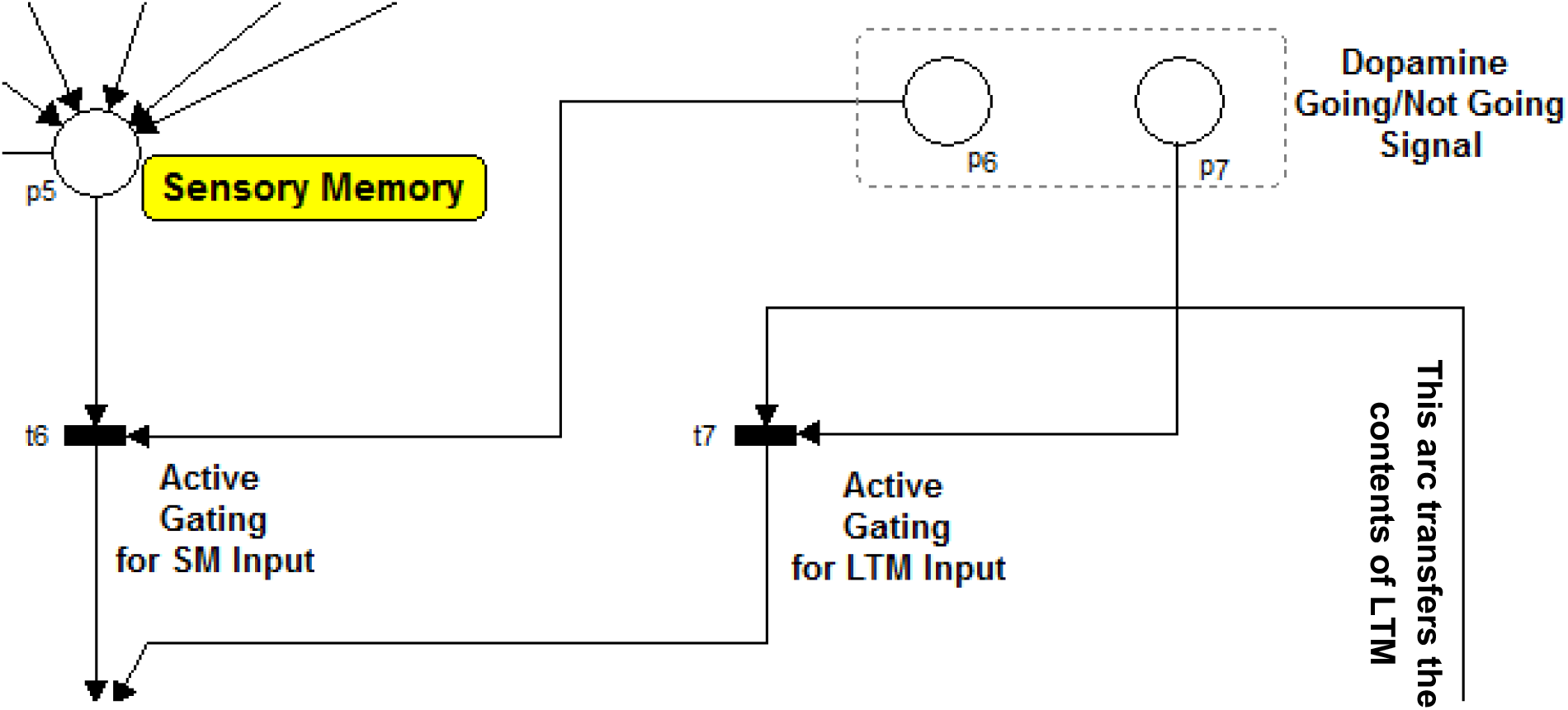
The PN modeling of active gating mechanism. Transitions t6 and t7 respectively control the pass of SM and LTM contents into WM. If a “Go” signal (a token) is sent to the place p6, then t6 can fire and let the SM content to flow into WM. Similarly, if a “Go” signal (a token) is sent to the place p7, then t7 can fire and let the LTM content to flow into WM. This model does not show how this “Go” signal is produced.

The inputs from SM should pass from the gate that has been modeled by an immediate transition t6. This transition has two input arcs and consequently, two input places (p5 and p6). According to the PN rules that have been described in section 2, transition t6 can fire whenever both its input places have at least one token. When SM receives an input stimulation, a token is placed in p6, and a half of the condition for firing the t6 is satisfied. The next part of the condition relates to a token in place p6. Without this token, t6 never allows SM to send its contents (tokens) to WM. When it detected that SM contains relevant information, dopaminergic neurons send the “Go” signal to p6. As discussed in the previous section, this detection is performed by the interaction between prefrontal cortex, basal ganglia, and WM (PBWM) through reinforcement learning (R. C. O’Reilly & Frank, 2006). The mechanisms of the mentioned detection and activation of dopaminergic neurons have not been modeled in our study. In the mathematical analysis and graphical animation in HPSim, we produced this signal (a token in place p6) manually. The same procedure is repeated for the inputs that come from LTM and request the permission to enter to WM. These inputs should pass transition t7. Opening or closing of this gate, t7, is controlled by the token (“Go” signal: that neurologically is related to Dopamine) in place p7.

In summary, when an incoming input from SM/LTM is relevant, a token (“Go” signal) is sent to place p6/p7. A token in these places can complete the mentioned condition and let the inputs to enter to WM, and we do not concern about how this token is produced in the proposed model. However, O’Reilly and Frank’s PBWM model (R. C. O’Reilly & Frank, 2006) can be added to the PN model to produce the “Go” signal in an appropriate time for the future development of the model.

Suppose that SM/LTM contents take the permission to flow into WM. The next concern of central executive part of working memory is to place the incoming data into the free cells of short-term storage buffer of WM that is discussed in the next subsection.

### 3.4. Filling the Short-term Storage Buffer Cells

In the proposed model, WM has two parts: a central executive function and short-term storage (STS) buffer. According to the previous researches, the storage buffer has a limited capacity (Hulme et al., 1995; Miller, 1956). Miller (1956) proposed a magical number 7±2 for the capacity of this STS buffer (Miller, 1956). In other words, we can say that our STS buffer has limited number of cells (7 ± 2), and each cell can store a unit of information that comes from SM or LTM for a short period (about 15 seconds) without considering “rehearsal mechanism.” In our model, we have considered five cells for STS buffer. The number of these cells can be increased or decreased in the proposed PN model. In Fig. 9, each of the places p9-p13 represents one of these cells. The inputs from SM and LTM that are allowed to pass the gates (the tokens in place p8) should be placed in STS buffer cells (places p9-p13) for further processing. In the PN model, corresponding to each of the STS buffer cells (p9-p13), we have considered another place that notifies the emptiness or fullness of that cell. According to Fig. 9, the places p19, p20, p21, p22, and p23 respectively notify the emptiness of the places p9, p10, p11, p12, and p13. For example, when place p19 has token, it means that place p9 (the first STS buffer cell) is empty and vice versa. Suppose that a unit of information (from SM or LTM) passes the active gate (t6 or t7 in Fig. 8). Therefore, there is a token in p8 that needs to be placed in one of the STS buffer cells. An immediate transition has been considered at the entrance of each STS buffer cell (t9-t13). These transitions act as a gate and prevent the overflow of the related cell. Each of these gates opens if its related cell is empty.

**Fig. 9.**
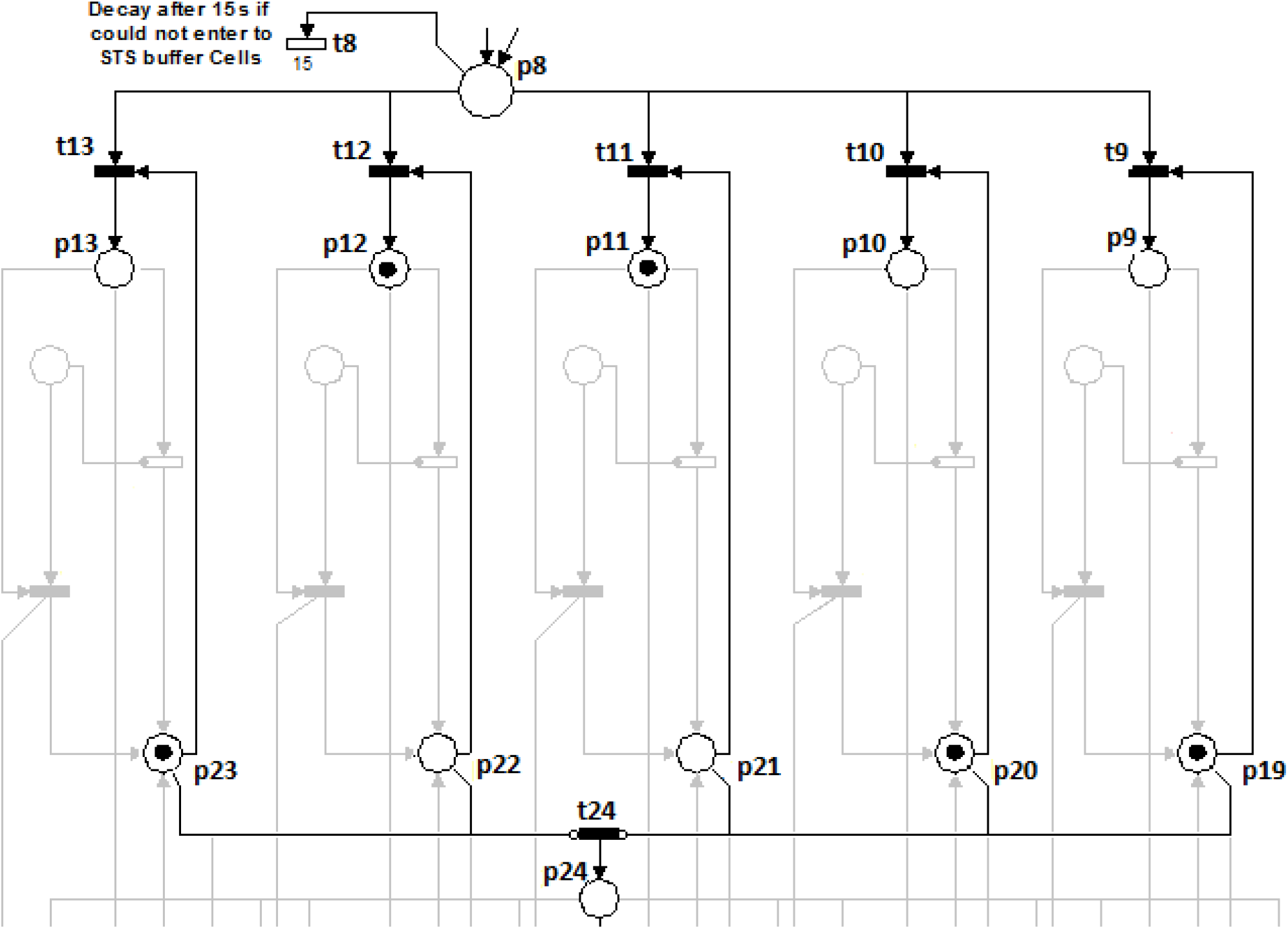
PN representation of filling the short-term storage buffer cells by incoming SM or LTM contents (tokens in p8). Five cells have been considered for STS buffer that is shown by p9-p13. When these cells (p9-p13) are full, the places (p19-p23) are empty and vice versa. Each of transition t9-t13 opens if its related cell is empty. This transition prevents the overflow of the buffer cells. If a token in p8 does not find the opportunity to flow into the cells, it decays from the time transition t8. Whenever all the cells become full, there is no token in p19-p23. These places are attached to the t24 by inhibitor arcs. Thus, t24 fires and send a token in p24, when all of its input places (p19-p23) have no token. Therefore, a token in p24 notifies the system that STS buffer is full.

For instance, suppose that there is a token in p8 that means a unit of information (from SM or LTM) has been detected as a relevant data and should be stored in one of the STS buffer cells. Fig. 9 shows that two cells (p11 and p12) are full. Therefore, the central executive unit should direct the incoming data to the free cells to avoid the overflow of full cells. This mechanism is implemented by the mentioned transitions t9-t13 and places p19-p23. According to Fig. 9, p21, and p22 have no token. It means that their corresponding buffer cells, p11 and p12, are full. Therefore, t11 and t12 cannot fire because one of their inputs places (p21 and p22) has no token. Consequently, the incoming data should flow into other cells (p9, p10, or p13). Suppose that it flows into p9. In the PN presentation, when t8 fires the token in p8 (an incoming data from SM/LTM) is sent to p9, and p19 loses its token that notifies the fullness of p9.

The next event occurs when all the cells are full, and relevant data passes the active gates and needs to be placed in one of the STS cells. Again, the central executive unit should select a mechanism to evacuate one of the cells immediately to avoid losing the data. In the PN presentation that has been shown in Fig. 9, if the evacuation procedure takes a long time, the token in p8 (relevant information from SM or LTM) decays through the time transition t8 or maybe overflow from p8 by the next incoming token (it is considered that p8 has limited capacity).

Whenever all the STS buffer cells (p9-p13) become full, there is no token in p19-p23. These places have been attached to t24 by inhibitor arcs. Thus, t24 fires and sends a token in p24 when all of its input places (p19-p23) have no token. Therefore, a token in p24 notifies that STS buffer is full.

By this notification, the central executive unit of the WM understands that a cell should be selected for evacuation upon the receiving of new data from active gates. Then, the content of the selected cell is updated with the new incoming input. This procedure is called “selective updating” in this paper.

### 3.5. Selective Updating Procedure

As it is mentioned before, in the current research, the term of “selective updating” refers to a procedure that central executive part of WM uses to select an STS buffer cell for updating its content with the new incoming information. A similar term was used in the study of O’Reilly and Frank (R. C. O’Reilly & Frank, 2006) and also in the study of Oberauer et al. (Oberauer et al., 2013). In those studies, this term (“selective updating”) refers to a mechanism that WM used for making selection between input stimuli and updating the content of STS buffer by the selected input. Indeed, in those studies, this mechanism was used to prevent the entrance of distracting stimuli to WM. For implementing this procedure, they used an active gating mechanism that was discussed in previous sections. In other words, in the mentioned studies, the term “selective updating” refers to the selection between input stimuli (from SM or LTM), and in our study, it refers to the selection between the content of STS buffer cells.

This selection would be critical. Suppose that the buffer cells are full and a new relevant incoming data arrives. If the central executive part makes an incorrect selection, the existent relevant information in a buffer cell will be updated by new relevant information. The previous information of the cell may be necessary to achieve a predefined goal. By an incorrect selection, WM loses required information for reaching to the goal of the task.

A common method for memory evacuation is the First-In-First-Out (FIFO) policy, which the data is removed from memory in the order that it has arrived. However, this policy is not appropriate in our work, because keeping the first arrived data may be necessary for completing a process. For example, when you want to multiply two numbers (e.g., 45 and 23) in your mind, the first entered data to WM is 45. This number should be kept in WM because it is necessary for completing the multiplication process. If the first number is removed from your STS buffer, the multiplication operation cannot be completed correctly.

Our first “selective updating” procedure is based on the last time that the content of each cell has been used during a task. The content of a cell that more time has passed from its last usage is supposed as the less important information.

To verify the proposed “selective updating” procedure, we have designed a test. Ten women whom their demographic characteristics are reported in Table 1 participated in this experiment.

**Table 1.** Demographic and Clinical Characteristics of Subjects

| Characteristic | Values |
| --- | --- |
| Gender | Female |
| Age | 27±3 |
| Number of the repetition of the test | one time |
| Duration of the test | 10 mins |
| Working memory span (visual digits) | 8±2 |

In this test, nine lists of two-digit numbers have been presented to participants visually at the rate of one number per two seconds. These lists are shown in Table 2.

**Table 2.**
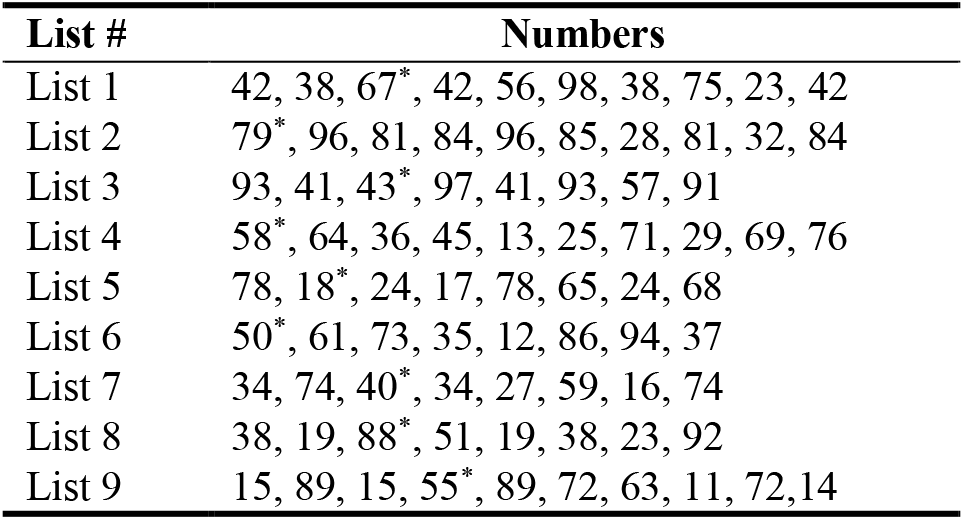
Nine lists of two-digit numbers. In each list, the number with a star indicates the number that more time has passed from its last observation.

Each list may contain repeated numbers. However, there are not two lists with a similar number. It can prevent from biasing by a number from a previous list. Participants were requested to memorize the numbers of that list without any concern about the order of the presentation (free recall). At the end of the list, different numbers were demonstrated on the screen. The contributor should immediately select the numbers that have observed during the previous stage (immediate free recall).

We hypothesized that the starred number in each list (the number that more time has passed from its last observation) might have less probability of recall concerning other numbers in the list. There were 10 participants in our experiment. If six subjects recalled a number in a list, the proportion of participants who have recalled the number would be 6/10 = 0.6. Therefore, the percentage of the probability of correctly recalling of that number is 60. Fig. 10 shows the probability of recall of the starred number (shown with square) and other numbers (shown with a cross) in each list.

**Fig. 10.**
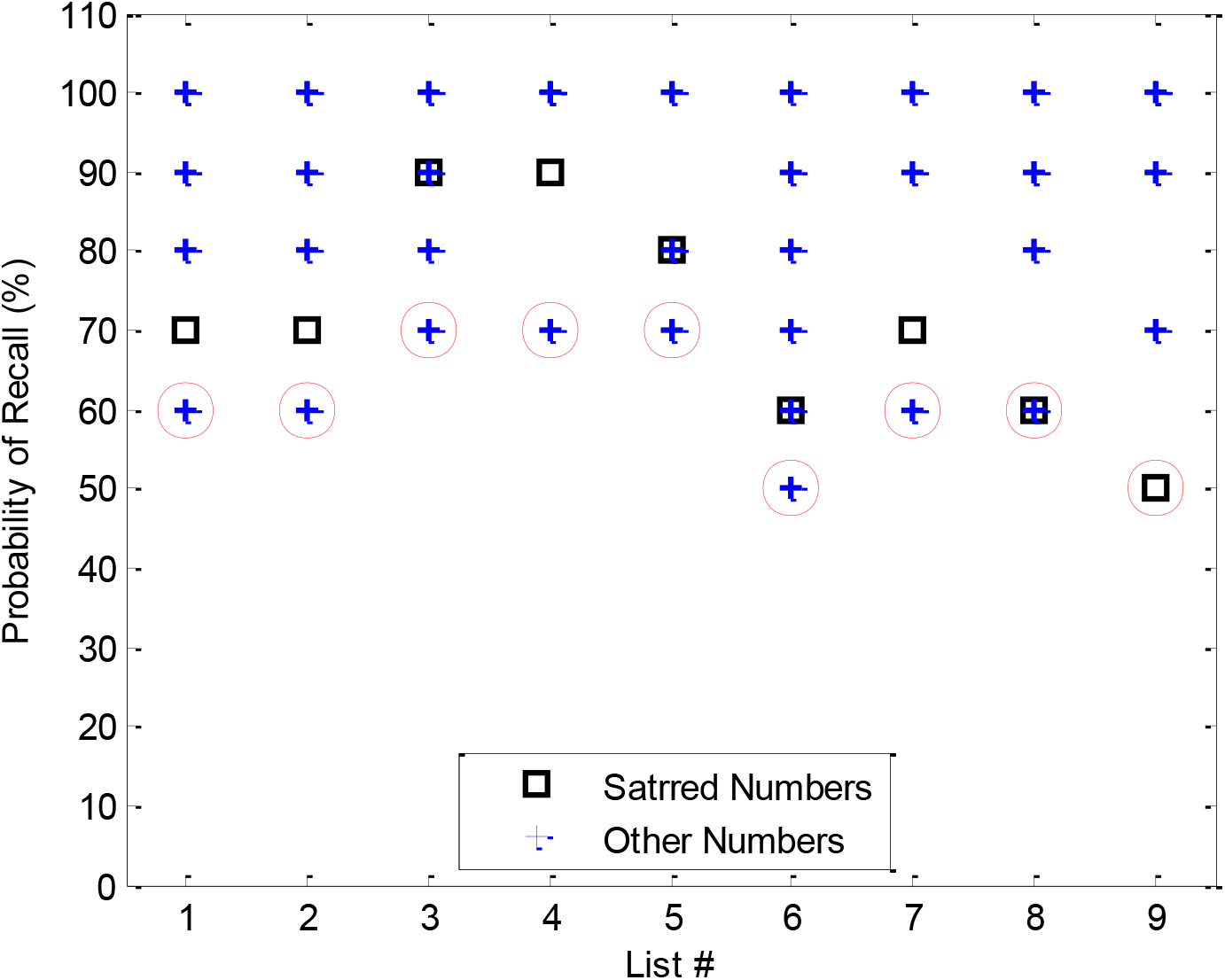
The probability of recall of the starred number and other numbers in each list. Squares: starred numbers, crosses: other numbers, and circles: indicate numbers that have the least probability of recall.

Fig. 10 shows that our hypothesis could not be true. Because there are numbers (indicated by circles in Fig. 10) that their recalling probability is less than the starred numbers (squares) in all lists except for 8th and 9th list. Closer examination of results revealed that probability of recall could relate to 1-the time that has passed from the last observation of an item and 2-the position of an item in a list. Studies on serial learning have shown that the items at the beginning and end of a list have a higher probability of recalling than items in the middle (Fig. 11) (Feigenbaum & Simon, 1962; Frensch, 1994). This phenomenon is called the “serial position effect.” The higher probability of recall of items at the beginning and the end of a list are respectively referred to as the “primacy” and “recency” effect (Li, 2010). The similar phenomenon was also observed in studies of free recalling (Murdock Jr, 1962).

**Fig. 11.**
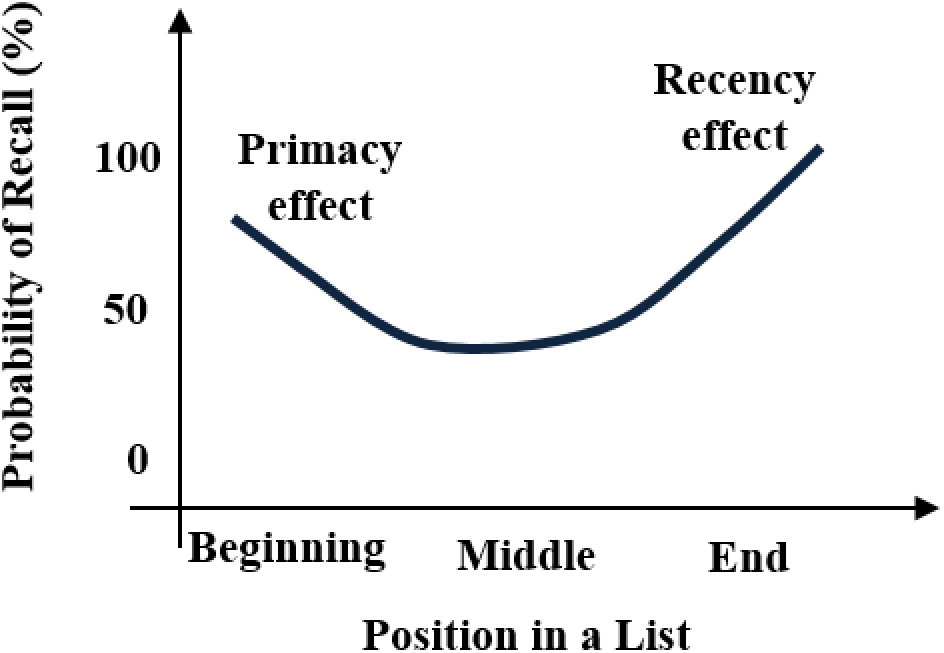
Serial position effect (in both free and serial recall)

According to the results of the experiment, we proposed a new procedure (shown in Fig. 12) to select a cell for updating its content with new incoming data.

**Fig. 12.**
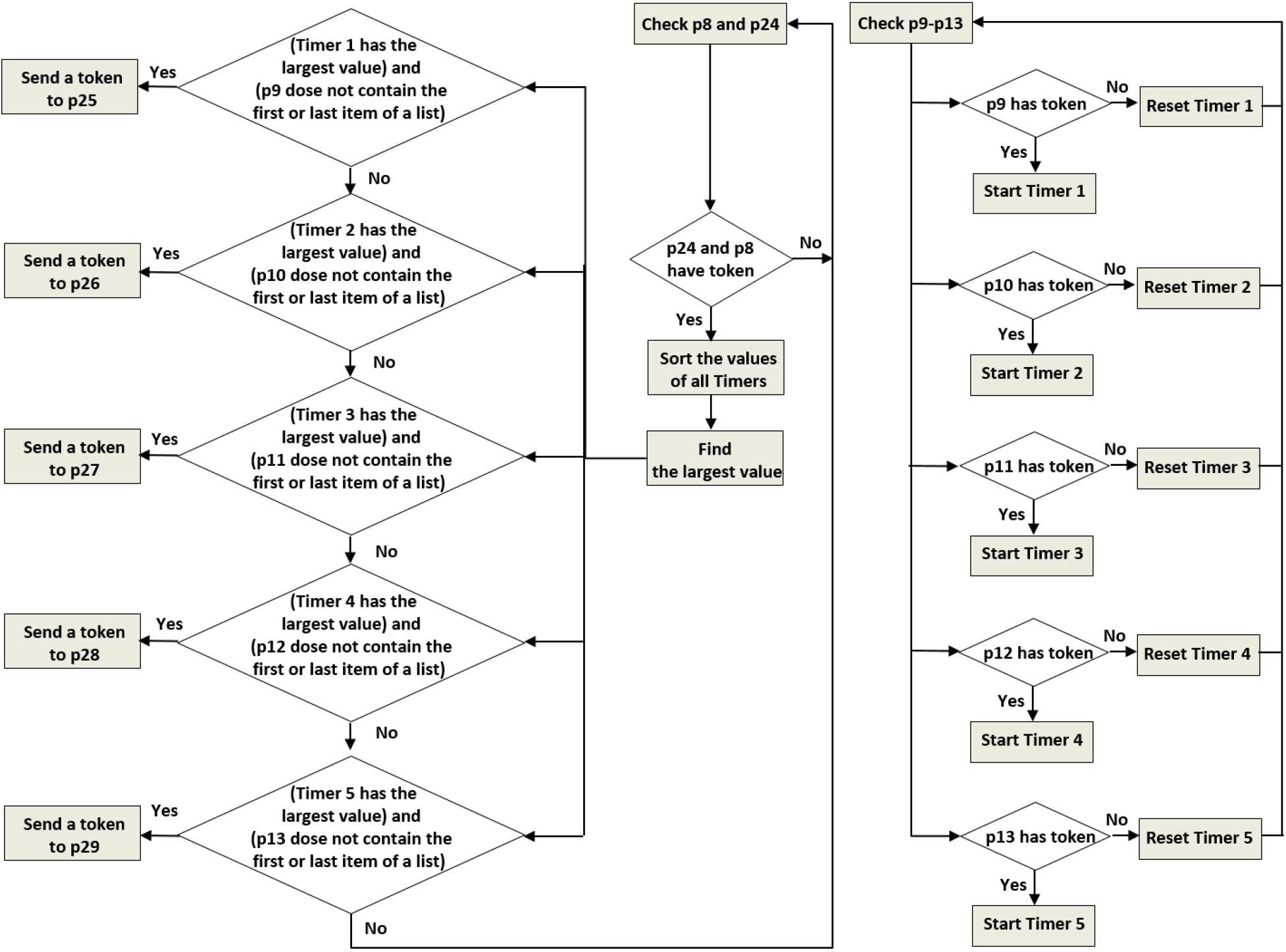
Selective updating procedure. A token is sent to the selected place.

According to the procedure shown in Fig. 12, the central executive part of the model decides to select a cell (place) for updating its content based on checking two conditions: 1-more time has passed from the last usage of the cell content, and 2-the cell content is not at the beginning of or the end of a list. For instance, suppose a new incoming relevant data that passes the active gate (a token in p8) and request the permission to flow into one of STS buffer cells. However, p24 has a token that indicates all STS buffer cells are full. A token in p24 notifies the central executive part to evacuate one of the cells. Again, suppose that more time has passed from the last usage of the content of p11 and it does not contain the first or the last item of a list. Therefore, the central executive part selects this place and sends a token to p27 to notify its decision. A token in this place (p27) shows that p11 has been chosen in the selective updating procedure. Similarly, a token in p25-p29 respectively shows the selection of p9-p13 (STS buffer cells) for updating. In this situation, all input places of t32 (p11: the selected STS buffer cell, p27: showing the selection of central executive part, and p24: showing the fullness of all buffer cells) have a token. Therefore, t32 fires and the previous token of p11 is updated by the new incoming token that comes from p8. The same procedure can occur for other cells. Fig. 13 demonstrates the overall PN presentation of the proposed model. All places and transitions of the net have been introduced in Table 3 and Table 4.

**Fig. 13.**
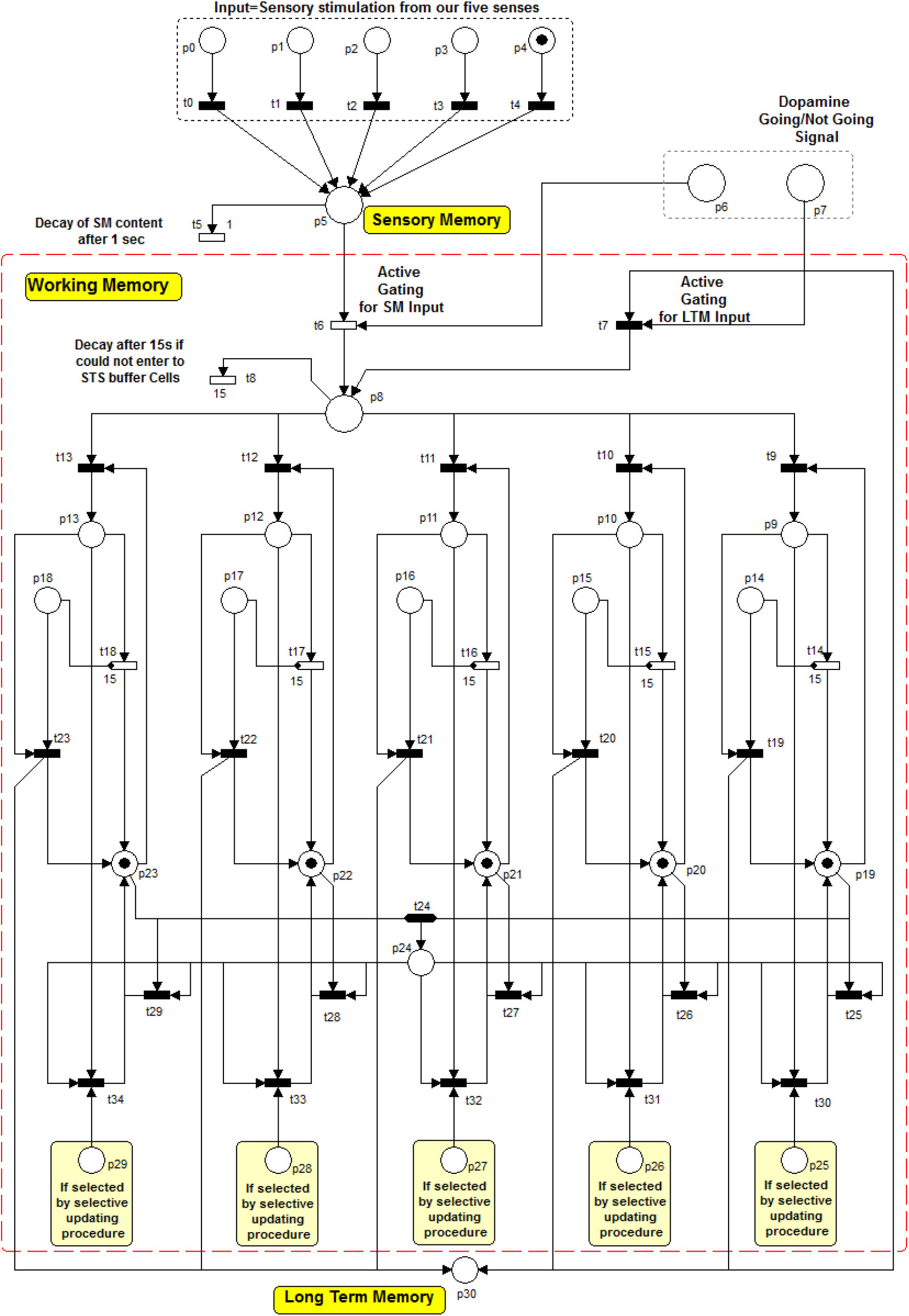
The overall PN presentation of the proposed model

**Table 3.** Interpretation of the places of the proposed model

| Place | Description |
| --- | --- |
| p0-p4 | Sensory inputs (Sight, hearing, smell, touch, taste) |
| p5 | SM |
| p6 | The command (“Go” signal) of opening the active gate to flow the data from SM to WM |
| p7 | The command (“Go” signal) of opening the active gate to flow the data from LTM to WM |
| p8 | The information that passes active gating and needs to store into an STS buffer cell |
| p9-p13 | STS buffer cells |
| p14-p18 | The required authority to send the content of an STS buffer cell to LTM |
| p19-p23 | STS buffer cell is empty |
| p24 | STS buffer cell is full |
| p25-p29 | Selected cell for evacuating by the “selective updating” method. |
| P30 | LTM |
| **SM:** Sensory Memory<br>**WM:** Working Memory<br>**LTM:** Long Term Memory<br>**STS:** Short Term Storage |  |

**Table 4.** Interpretation of transitions of the proposed model

| Transition | Description |
| --- | --- |
| t0-t4 | Transmission of sensory input to SM |
| t5<sup>*1</sup> | The decay of SM content |
| t6 | The active gate that controls the entrance of SM content to STS buffer cell |
| t7 | The active gate that controls the entrance of LTM content to STS buffer cell |
| t8<sup>*2</sup> | The decay of information that passes the active gates but could not flow into STS buffer cell |
| t9-t13 | Transmission of the information that passes the active gate to STS buffer cells |
| t14-t18<sup>*3</sup> | The decay of the content of STS buffer cells |
| t19-t23 | Transmission of information from STS buffer cells to LTM |
| t24 | Sending the notification of the fullness of STS buffer |
| t25-t29 | Eliminate the token of p24 whenever one of the buffer cells becomes empty |
| t30-t34 | Evacuating the selected STS buffer cell by the “selective updating” procedure |

<sup>\*</sup>Timed transition
<sup>1</sup>Time delay of t5=500 milliseconds
<sup>2</sup>Time delay of t8=15 seconds
<sup>3</sup>Time delay of t14-t18=15 seconds

The other possible method to evacuate an STS buffer cell is to send its content into the LTM, which has been described in the following subsection.

### 3.6. The evacuation of STS buffer cells

In the proposed model, three scenarios have been considered to evacuate an STS buffer cell. One scenario was discussed in the previous section. The second scenario is to send the information from STS buffer to LTM regardless of fullness or emptiness of STS buffer cells. Such a condition can occur when the content of a cell can find the required authority to encode into the LTM. For instance, suppose that the content of p11 has the authority to encode into LTM. If p11 has such authority, a token is sent into p16. This place, p16, is considered for holding the permission of sending the content of p11 into LTM. In the current version of the proposed model, we do not concern about where and how this authority is identified. According to Fig. 13, when p11 and p16 have token, t21 fires and sends the content (token) of p11 to LTM that has been presented by p30 in the proposed model, and send a token to p21 to notify the emptiness of p11. This event can occur in other cells (p9-p13) through receiving the authority in places p14-p18 and firing the transitions t19-t23. However, what will happen for the content of an STS buffer cell if it does not receive the authority to flow into LTM. As it is mentioned in the previous section, STS buffer can hold the incoming data for a short period, about 15 seconds without any rehearsal. After this short period, the third scenario happens to evacuate the buffer.

We have not considered the rehearsal mechanism in our proposed model. Therefore, when a unit of information enters into a buffer cell and cannot flow into LTM or is not updated by the selective updating procedure, it decays automatically after about 15 seconds. According to Fig. 13, p11 (one of the cells of STS buffer cells) has one input and three output arcs. The input arc transfers the information from SM or LTM into this cell (p11). The first output arc transfers the content of p11 into LTM (p30), the second arc evacuates the content of p11 through “selective updating” procedure, and the third arc has been considered to decay the content of p11 after about 15 seconds. This arc has been attached to a timed transition (t16). The time delay of this transition is tuned on 15 seconds. As can be seen in Fig. 13, an inhibitor arc has attached from place p16 (the place that holds the permission of sending the content of p11 into LTM) to the timed transition t16. This inhibitor arc has been considered to resolve a possible conflict. Suppose that 15 seconds pass from the time of the entrance of information to p11. Therefore, t16 is ready for firing. At the same time, the content of p11 finds the authority to flow into LTM (a token is sent to p16). In this situation, both t16 and t21 are ready for firing simultaneously. Thus a conflict occurs between t6 and t21 to use the content of p11. If t16 is allowed to fire, WM lose relevant data that may require it to complete a predefined task. To prevent the firing of t16, an inhibitor arc from p16 has been attached to t16. Since p16 has token, t16 cannot fire.

Same arcs have been considered for other cells of STS buffer (i.e., p9-p13). Therefore, each of STS buffer cells can evacuate through one of the mentioned three scenarios. Suppose that STS buffer is full then central executive part of WM sends a notification (i.e., a token is sent to p24). Then assume that p11 is evacuated through one on these scenarios. The evacuation is notified by sending a token in p21. As soon as one of the STS buffer cells (here p11) becomes empty, central executive part of WM should remove its notification about the fullness of the buffer. In other word, if there is a token in p24, it should be removed. Fig. 13 shows that the firing of t27 leads to the elimination of the mentioned notification (i.e., token in p24). This transition, t27, has two input places. One of them is p24 that has a token whenever the buffer is full. The other one is p21 that has token whenever p11 (one of the buffer cells) becomes empty. Such a transition has been considered for other STS buffer cells (i.e., t25-t29 respectively corresponds to p9-p13).

## 4. Comparison of the proposed model with other computational models of working memory

## 5. The Analysis Result of the Proposed Model: The Prediction of Inattention Symptoms Causes

In previous sections, different parts of the proposed model were described. In this section, it is tried to investigate the proposed model more precisely to predict the possible causes of inattention symptoms that have been listed below (Nigg, 2006):

- Easily distracting by the environmental noise
- Difficulty in sustained attention
- Excessive daydreaming during the work
- Inability to finish work and excessive switching from on a task to another one
- Re-reading a text because of difficulty in focus maintenance

People with such symptoms usually make more errors than normal people during different attentional tests (Lundervold et al., 2011). Therefore, they are labeled as inattentive subjects. As a result of different studies, it has been confirmed that WM capacity has an important role in controlling attention (Cowan et al., 2005; Kane & Engle, 2003). In these studies, it was claimed that low WM capacity could lead to neglect the goal of the test before completing it and consequently, could increase the performance error (Kane & Engle, 2003). In other words, low WM capacity causes the overflow of buffers, losing the goal, and thereby appearing inattentive symptoms.

According to the proposed PN model, it can be seen that moreover than the low WM capacity (decreasing the number of STS buffer cells), the following reasons may also lead to loss of useful information:

1. SM is one of the sources that provide required relevant information of performing a task for WM. If WM receives this information incompletely, the task cannot be executed correctly and leads to performance error. According to the PN proposed model (Fig. 13), suppose that the processing speed of WM is much less than the speed of receiving information from SM. The STS buffer of WM (p9-p13) becomes full by the incoming information from SM. The next incoming stimuli should wait until the evacuation of STS buffer cells. The evacuation procedure is performed by the central executive part of the WM. If this part operates slowly, the incoming information overflows (SM has limited capacity, about 12 units of information (Sperling, 1963)). Therefore, WM loses some of the required information to complete the task correctly. This result leaves one question: “How fast the stimuli input should apply to avoid this problem?” A delay about T1 was added to the block of “selective updating procedure,” and the speed of incoming stimuli into SM is tuned on T2 to simulate such a situation in the proposed PN model. Then, we ran the model and tried to observe this problem (i.e., overflow of SM (p5)) by changing the ratio between T1 and T2. In this simulation, the maximum capacity of SM was set on 12 (according to (Sperling, 1963)). The result will change by increasing or decreasing this capacity. For simplicity of simulation, we tuned T1 on 1ms, and changes were done on the value of T2. Table 5 shows the result of this simulation. According to Table 5, it can be seen that when the inter-stimuli interval (T1) is shorter than 2ms, there is no overflow. In this condition, the optimum speed of delivering input stimuli can vary from one subject to another subject because the speed of WM processing is different between subjects. Table 5 shows that decreasing the WM processing speed (i.e., increasing T2) leads to the sooner occurrence of overflow. It has been reported that people with attention disorder can concentrate on a task shorter than normal subjects (Barkley, 1997). It can be suggested that the in people with attention deficit disorder, central executive part of WM may perform the selective updating procedure slower than normal ones. The mentioned overflow may also lead to losing some relevant information to complete a task. Therefore, the subject may prefer to jump to do another task (task switching symptom in people with attention disorder (Nigg, 2006)). Another interesting result that is observed in the seventh row of Table 5 is that by little increasing of the inter-stimuli interval, the low speed of central executive part of WM can be compensated (see Fig. 14). In the first row, T1=1ms and T2=10ms, and the first overflow in SM (p5) occurred after 120ms. In the last row by increasing T1 just for 2ms (T1=3ms and T2=10ms), the overflow problem was eliminated.
2. In the previous part, the effect of the delay of the central executive part of WM in evacuating STS buffer cells was investigated. In this part, it is assumed that SM sends units of information into STS buffer with delay. Suppose that you are reading a newspaper. To understand the concept of a sentence, WM needs to put the words beside each other. If your SM sends the information of these words into STS buffer very slowly, the decay of previous words is possible in STS buffer. Therefore, WM may lose some of the words that are necessary to find the concept of the sentence. This situation was simulated by adding some delay to transition t6. Thus, the content of SM (p5) is sent to the STS buffer of WM with some delay, and consequently, the decay of the content of STS buffer cells (p9-p13) may occur. Table 6 shows the flow of tokens (units of information) in STS buffer cells by considering the mentioned delay. According to Table 6, it can be seen that in step 3, the content of p9 has decayed and has been rewritten by new information in step 4. Therefore, WM has lost some of the required information in step3. It can be suggested that excessive re-reading symptom of people with an attention disorder may be due to this problem. In that, they have to re-read the sentence to find the missed words. The result of the simulation in Table 5 showed that increasing the time between delivering input stimuli could compensate for the delay of the central executive part. But, how the delay of sending data from SM into WM can be compensated? It seems that in people with such a problem, the proposed “rehearsal loop” of Baddeley (Fig. 4) may have a more important role. The rehearsal loop helps WM to keep the information in STS buffer cells for a longer period. Therefore, training these people to use their “rehearsal loop” mechanism can help them to compensate for the delay of sending information from SM to WM (see Fig. 15).
3. As it is discussed in the previous sections, selective gates should be opened in an appropriate time to allow the relevant information flows into STS buffer of WM from SM or LTM. SM can hold information for a short period. Suppose that the “Go” signal (token in place p6) is received late (takes time larger than one second). Therefore, before firing of t6 (active gate of SM), the content of SM decays. It can be considered as the second root causes of re-reading symptoms in people with an attention disorder. In addition to the decay of relevant information from SM, the entrance of irrelevant information is also possible. For example, a unit of relevant information requests the permission to flow into STS buffer of WM at the time k1, and it sends its request and waits for the permission of passing the active gate. This permission (“Go” signal) is sent to the active gate at time k1+1100ms. The delay between sending the request and receiving the permission of passing the gate is more than one second. Therefore, this relevant information decay from SM and the permission of opening the gate is useless. However, the gate is opened by this permission at the time k1+1100ms. If SM receives a unit of irrelevant information at the same time (i.e., k1+1100ms), it can easily pass from the opened gate and occupies one of the STS buffer cells of WM. This problem may lead to distraction. Fig. 16 shows this problem graphically.
4. As mentioned before, an inhibitor arc has been connected from p14-p18 to t14-t18 to solve a possible conflict. This conflict occurs when the permission of sending the content of an STS buffer cell into LTM is synchronized with the time of the automatic decay. The mentioned inhibitor arcs prevent the decay of information in such situations. Thus, if the mentioned permission is received with a very short delay after the time of automatic delay, the inhibitor arc became ineffective, and the information decays form WM. When you have to sustain your attention on work for some time, your WM should have appropriate communication with LTM to keep the sequential stages of the work in your mind. Due to the limited capacity of WM, it sends some of the information into the LTM and recalls them at any time that is needed. If the permission of sending data to LTM receives with an unexpected delay, some stages of the work may lose, and you cannot finish the work correctly.
5. Over-focusing problem is another symptom that has rarely been observed in people with attention deficit disorder. In this problem, people have trouble to shift their attention from one task to another one. This symptom is often called Obsessive-Compulsive Disorder (OCD) (Masi et al., 2006). People with this symptom are over-attentive instead of inattentive. The proposed PN model can also show some reasons for the source of this symptom. If the pathways that have been considered to evacuate the STS buffer cells damage (as a result of brain injury, congenital disabilities or some chemical drug consumption), the content of WM cannot be updated with new information, therefore, “working” on the previous content of “memory” cells is continued, and the new task or thought cannot be started. The same result may occur if the parts that send the command of opening these pathways do not work correctly (in the PN model, the command is a token in p25-p29 and p14-p18). Anatomically, these parts are associated with the frontal regions of the brain. Previous brain studies showed that damages of the frontal lobe might lead to the impairment of WM and attention system function (Kates et al., 2002; Kimberg & Farah, 1993).

**Fig. 14.**
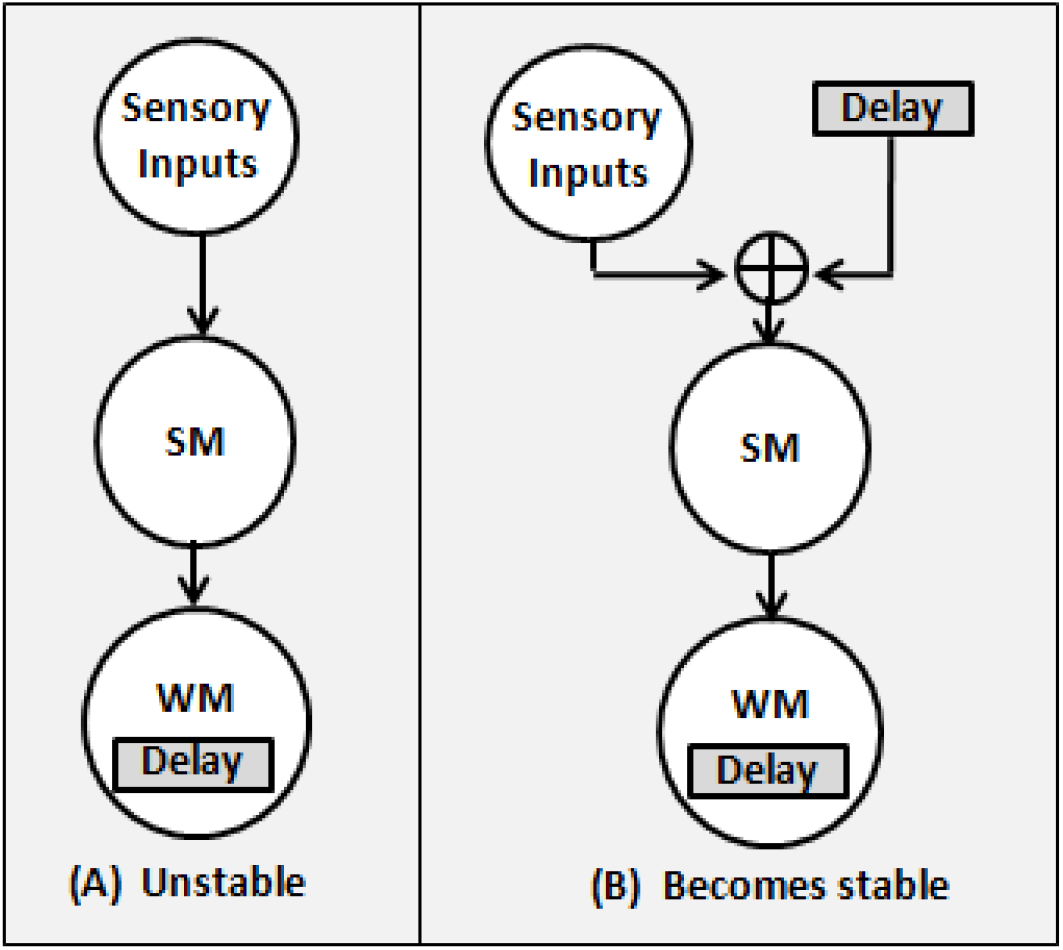
(A) SM overflows and becomes unstable because of the delay of the central executive part of WM to evacuate STS buffer cells. (B) Compensating the WM delay by adding a delay to the time between delivering different input stimuli. It avoids the overflow of SM, and the system becomes stable.

**Fig. 15.**
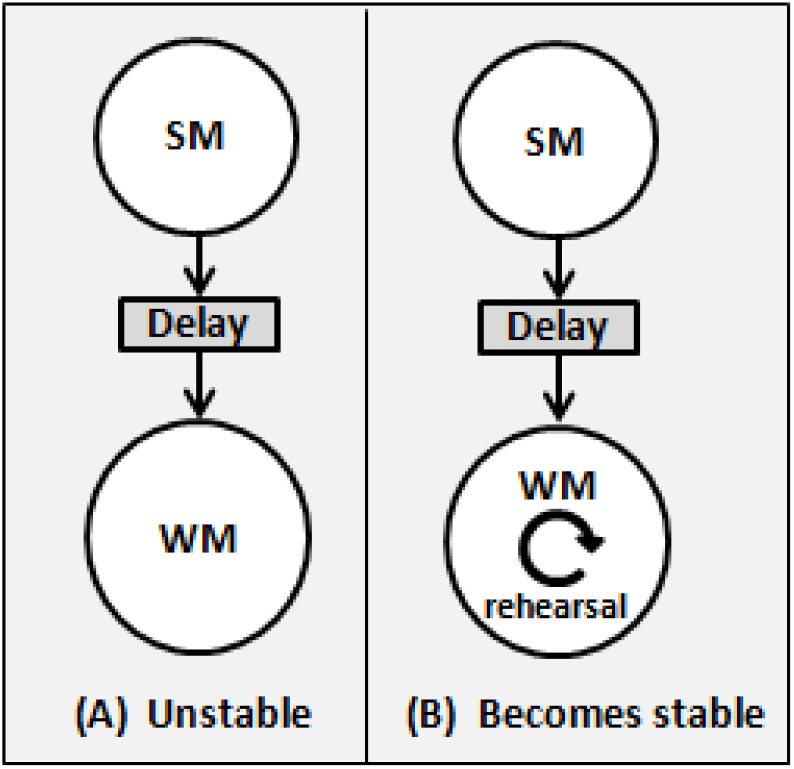
(A) Some of the information decay from STS buffer because of a delay in sending required information from SM to WM. Therefore, the system becomes unstable. (B) By the rehearsal mechanism, WM can hold information for more time, and the mentioned delay is compensated, and the system becomes stable.

**Fig. 16.**
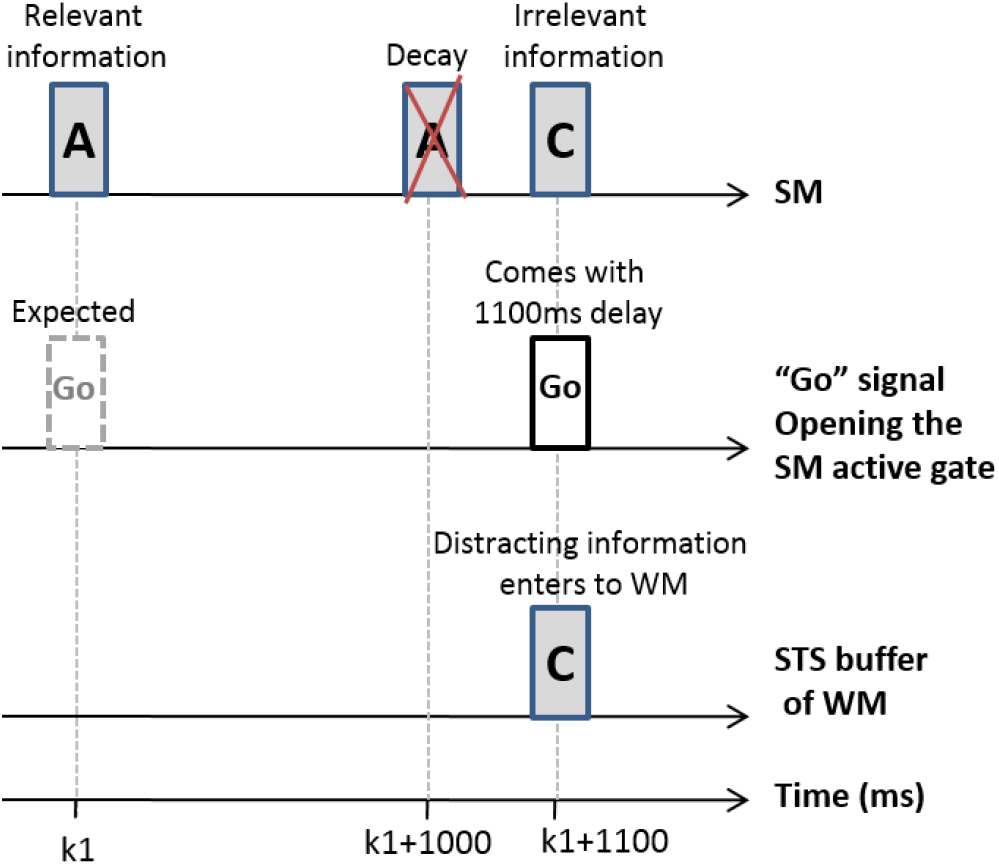
“A” is considered as relevant, and “C” is considered as irrelevant (distracting) information. “A” comes to SM at the time k1. Therefore, a “Go” signal (a token in p6) is expected at the time k1. “Go” signal comes with 1100 ms delay. “A” decays after 1000 ms. Thus WM loses a unit of relevant information. A unit of irrelevant information (“C”) comes to SM at the time k1+1100ms. “Go” signal comes with a delay at the same time. Therefore, the gate is opened, and “C” passes the gate, flows into WM, and may lead to distraction.

**Table 5.** Investigating the effect of the time the central executive part required to evacuate an STS buffer cell using “selective updating” procedure (T2) concerning the inter-stimuli interval (T1)

|  | **T1** | **T2** | **The elapsed time that the first overflow in SM (p5) occurred** |
| --- | --- | --- | --- |
| 1 | 1ms | 10ms | 120ms |
| 2 | 1ms | 8ms | 130ms |
| 3 | 1ms | 5ms | 149ms |
| 4 | 1ms | 3ms | 205ms |
| 5 | 1ms | 2ms | 325ms |
| 6 | 1ms | 1ms | No Overflow |
| 7 | 3ms | 10ms | No Overflow |

**Table 6.**
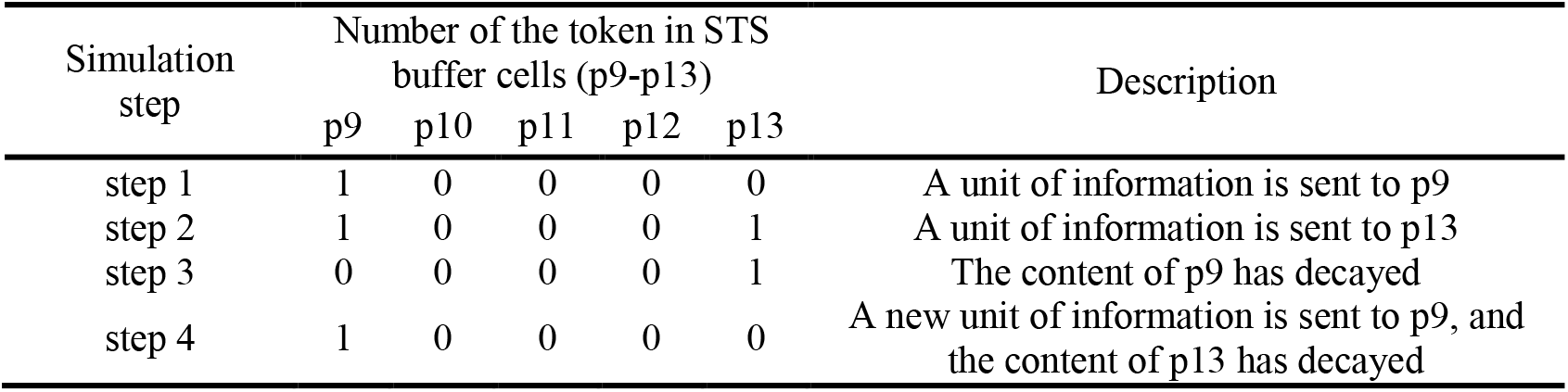
The flow of tokens (units of information) in STS buffer cells by adding some delay to transition t6 to simulate the SM transmission delay.

| Simulation step | Number of the token in STS buffer cells (p9-p13) |  |  |  |  | Description |
| --- | --- | --- | --- | --- | --- | --- |
|  | p9 | p10 | p11 | p12 | p13 |  |
| step 1 | 1 | 0 | 0 | 0 | 0 | A unit of information is sent to p9 |
| step 2 | 1 | 0 | 0 | 0 | 1 | A unit of information is sent to p13 |
| step 3 | 0 | 0 | 0 | 0 | 1 | The content of p9 has decayed |
| step 4 | 1 | 0 | 0 | 0 | 0 | A new unit of information is sent to p9, and the content of p13 has decayed |

These suggestions may be valuable to find the possible defects of brain components that may lead to attention disorder symptoms. They can be used to propose new treatments to improve the inattentive behavior of people with ADD.

## 6. Conclusion

In this study, a PN model was proposed to show the flow of information from SM and LTM into WM. In this model, WM was divided into two parts: STS buffer to store a limited amount of information (five items) for a short time (15 seconds without any rehearsal) and an executive central part of controlling the flow of information between different parts (SM, LTM, STS buffer). The basis of this model was formed according to the biological evidence that was reported in the previous studies. However, a new “selective updating” mechanism was also proposed. Central executive part of WM used this mechanism to update the less important contents of STS buffer with new incoming information. The proposed PN model showed the importance of the effect of delays in the flow of information between different parts of the model (e.g., SM, LTM, WM (STS buffer + Central executive)).

It was also observed that changing the timing of different procedure affected the performance of the system and led to different symptoms of inattention.

For future works, we have decided to add the mechanism that detects when “Go” signals should be sent to open the active gates. This development can be implemented by PN presentation of the O’Reilly and Frank (2006) computational model that works based on reinforcement learning. The other mechanism that can be added into the model is the Baddely’s rehearsal mechanism. These extensions can be valuable to find out other possible causes of inattention symptoms.

## Notes

**Declarations of interest:** none

### Competing Interest Statement

The authors have declared no competing interest.

## References

Arias-Carrión, Ó., & Pöppel, E. (2007). Dopamine, learning, and reward-seeking behavior. J Acta neurobiologiae experimentalis.

Aries, R. J., Groot, W., & van den Brink, H. M. (2015). Improving reasoning skills in secondary history education by working memory training. J British Educational Research Journal, 41(2), 210–228.

Ashby, F. G., Ell, S. W., Valentin, V. V., & Casale, M. B. (2005). FROST: A distributed neurocomputational model of working memory maintenance. Journal of Cognitive Neuroscience, 17(11), 1728–1743.

Atkinson, R. C., & Shiffrin, R. M. (1968). Human Memory: A Proposed System and its Control Processes. In K. W. Spence & J. T. Spence (Eds.), Psychology of Learning and Motivation (pp. 89–195): Academic Press.

Awh, E., Vogel, E. K., & Oh, S.-H. (2006). Interactions between attention and working memory. J Neuroscience, 139(1), 201–208.

Baddeley, A. (2000). The episodic buffer: a new component of working memory? Trends in Cognitive Sciences, 4(11), 417–423.

Baddeley, A. (2003). Working Memory: Looking Back and Looking Forward. Nature Reviews Neuroscience, 4(10), 829–839.

Badre, D. (2012). Opening the gate to working memory. Proceedings of the National Academy of Sciences of the United States of America, 109(49), 19878–19879.

Baghdadi, G., Towhidkhah, F., & Rostami, R. (2017). An electrophysiological model of working memory performance. Cognitive Systems Research, 45, 1–16.

Barkley, R. A. (1997). Behavioral inhibition, sustained attention, and executive functions: Constructing a unifying theory of ADHD. Psychological Bulletin, 121(1), 65–94.

Bayerl, M., Dielentheis, T. F., Vucurevic, G., Gesierich, T., Vogel, F., Fehr, C., et al. (2010). Disturbed brain activation during a working memory task in drug-naive adult patients with ADHD. J Neuroreport, 21(6), 442–446.

Bays, P. M. (2018). Reassessing the Evidence for Capacity Limits in Neural Signals Related to Working Memory. Cerebral Cortex, 28(4), 1432–1438.

Biederman, J. (1998). Attention-deficit/hyperactivity disorder: a life-span perspective. The Journal of clinical psychiatry, 59, 4–16.

Blätke, M. A., Heiner, M., & Marwan, W. (2011). Tutorial - Petri Nets in Systems Biology. Technical Report.

Borella, E., Carretti, B., Riboldi, F., & De Beni, R. (2010). Working memory training in older adults: Evidence of transfer and maintenance effects. Psychology and Aging, 25(4), 767–778.

Bouyer, P., Haddad, S., & Reynier, P.-A. (2008). Timed Petri nets and timed automata: On the discriminating power of zeno sequences. Information and Computation, 206(1), 73–107.

Brown, G. D., Hulme, C., & Preece, T. (2000). Oscillator-Based Memory for Serial Order. Psychological Review, 107(1), 127–181.

Burgess, N., & Hitch, G. J. (1992). Toward a Network Model of the Articulatory Loop. Journal of Memory and Language, 31(4), 729–760.

Chun, M. M., & Turk-Browne, N. B. (2007). Interactions between attention and memory. Curr Opin Neurobiol, 17(2), 177–184.

Cowan, N. (2008). What are the differences between long-term, short-term, and working memory? J Progress in brain research, 16(9), 323–338.

Cowan, N., Elliott, E. M., Scott Saults, J., Morey, C. C., Mattox, S., Hismjatullina, A., et al. (2005). On the capacity of attention: Its estimation and its role in working memory and cognitive aptitudes. Cognitive Psychology, 51(1), 42–100.

Damadi, S. M. S., Doustmohammadi, A., & Afshar, A. (1989). Decentralized supervisory control of discrete event systems using Petri nets. Amirkabir university of Technology of Iran.

Dowson, J., McLean, A., Bazanis, E., Toone, B., Young, S., Robbins, T., et al. (2004). Impaired spatial working memory in adults with attention-deficit/hyperactivity disorder: comparisons with performance in adults with borderline personality disorder and in control subjects. J Acta Psychiatrica Scandinavica, 110(1), 45–54.

Engle, R. W., Kane, M. J., & Tuholski, S. W. (1999). Individual differences in working memory capacity and what they tell us about controlled attention, general fluid intelligence, and functions of the prefrontal cortex. In Models of working memory: Mechanisms of active maintenance and executive control. (pp. 102–134). New York, NY, US: Cambridge University Press.

Fassbender, C., Schweitzer, J. B., Cortes, C. R., Tagamets, M. A., Windsor, T. A., Reeves, G. M., et al. (2011). Working memory in attention deficit/hyperactivity disorder is characterized by a lack of specialization of brain function. JPloS one, 6(11), e27240.

Feigenbaum, E. A., & Simon, H. A. (1962). A Theory of the Serial Position Effect. British Journal of Psychology, 53(3), 307–320.

Fougnie, D. (2008). The relationship between attention and working memory. New research on short-term memory, 1, 45.

Frank, M. J. (2005). Dynamic Dopamine Modulation in the Basal Ganglia: A Neurocomputational Account of Cognitive Deficits in Medicated and Nonmedicated Parkinsonism. J. Cognitive Neuroscience, 17(1), 51–72.

Frank, M. J., Loughry, B., & O’Reilly, R. C. J. C., Affective,. (2001). Interactions between frontal cortex and basal ganglia in working memory: A computational model. J Cognitive, Affective, Behavioral Neuroscience, 1(2), 137–160.

Frensch, P. (1994). Composition During Serial Learning: A Serial Position Effect (Vol. 20).

Friedman, D., & Johnson Jr., R. (2000). Event-related potential (ERP) studies of memory encoding and retrieval: A selective review. 51(1), 6–28.

Gruber, A. J., Dayan, P., Gutkin, B. S., & Solla, S. A. (2006). Dopamine modulation in the basal ganglia locks the gate to working memory. Journal of Computational Neuroscience, 20(2), 153.

Henson, R. N. (1998). Short-Term Memory for Serial Order: The Start-End Model. COGNITIVE PSYCHOLOGY, 36, 73–137.

Hrz, B., & Zhou, M. C. (2007). Modeling and Control of Discrete-event Dynamic Systems: with Petri Nets and Other Tools: Springer Publishing Company, Incorporated.

Hulme, C., Roodenrys, S,. Brown, G., & Mercer, R. (1995). The role of long-term memory mechanisms in memory span. 86(4), 527–536.

Jarrold, C., Baddeley, A. D., & Phillips, C. (1999). Down syndrome and the phonological loop: The evidence for, and importance of, a specific verbal short-term memory deficit. Down Syndrome: Research & Practice, 6(2), 61–75.

Joo, J., Kim, N., Wysk, R., Rothrock, L., Son, Y.-J., Oh, Y. G., et al. (2013). Agent-based simulation of affordance-based human behaviors in emergency evacuation (Vol. 32).

Kane, M. J., & Engle, R. W. (2003). Working-memory capacity and the control of attention: The contributions of goal neglect, response competition, and task set to Stroop interference. Journal of Experimental Psychology: General, 132(1), 47–70.

Kates, W. R., Frederikse, M., Mostofsky, S. H., Folley, B. S., Cooper, K., Mazur-Hopkins, P., et al. (2002). MRI parcellation of the frontal lobe in boys with attention deficit hyperactivity disorder or Tourette syndrome. J Psychiatry Research: Neuroimaging, 116(1-2), 63–81.

Kimberg, D. Y., & Farah, M. J. (1993). A unified account of cognitive impairments following frontal lobe damage: The role of working memory in complex, organized behavior. J Journal of Experimental Psychology: General, 122(4), 411.

Klados, M. A., Simos, P., Micheloyannis, S., Margulies, D., & Bamidis, P. D. (2015). ERP measures of math anxiety: how math anxiety affects working memory and mental calculation tasks? J Frontiers in behavioral neuroscience, 9, 282.

Klingberg, T., Fernell, E., Olesen, P. J,. Johnson, M., Gustafsson, P., Dahlström, K., et al. (2005). Computerized training of working memory in children with ADHD-a randomized, controlled trial. Journal of the American Academy of Child Adolescent Psychiatry, 44(2), 177–186.

Klingberg, T., Forssberg, H., & Westerberg, H. (2002). Training of working memory in children with ADHD. Journal of clinical experimental neuropsychology, 24(6), 781–791.

Kobel, M., Bechtel, N., Specht, K., Klarhöfer, M., Weber, P., Scheffler, K., et al. (2010). Structural and functional imaging approaches in attention deficit/hyperactivity disorder: does the temporal lobe play a key role? J Psychiatry Research: Neuroimaging, 183(3), 230–236.

Kobel, M., Bechtel, N., Weber, P., Specht, K., Klarhöfer, M., Scheffler, K., et al. (2009). Effects of methylphenidate on working memory functioning in children with attention deficit/hyperactivity disorder. J European Journal of Paediatric Neurology, 13(6), 516–523.

Kofler, M., Rapport, M., Bolden, J., Sarver, D., & Raiker, J. (2009). ADHD and Working Memory: The Impact of Central Executive Deficits and Exceeding Storage/Rehearsal Capacity on Observed Inattentive Behavior. Journal of abnormal child psychology, 38(2), 149–161.

Kouzehgar, M., Badamchizadeh, M. A., & Khanmohammadi, S. (2013). Fuzzy Petri Nets for Human Behavior Verification and Validation. arXiv preprint arXiv:1303, 1247.

Kuhl, B. A., & Chun, M. (2014). Memory and attention. In The Oxford handbook of attention.

Lewandowsky, S., & Farrell, S. (2005). Computational models of working memory. Encyclopedia of Cognitive Science, 1–10.

Li, C. (2010). Primacy effect or recency effect? A long-term memory test of Super Bowl commercials. Journal of Consumer Behaviour: An International Research Review 9(1), 32–44.

Lofthouse, N., Arnold, L. E., Hersch, S., Hurt, E., & DeBeus, R. (2011). A Review of Neurofeedback Treatment for Pediatric ADHD. Journal of Attention Disorders, 16(5), 351–372.

Lundervold, A. J., Adolfsdottir, S., Halleland, H., Halmøy, A., Plessen, K., & Haavik, J. (2011) Attention Network Test in adults with ADHD - the impact of affective fluctuations. J Behavioral Brain Functions, 7(1), 27.

Martinussen, R., Hayden, J., Hogg-Johnson, S., & Tannock, R. (2005). A meta-analysis of working memory impairments in children with attention-deficit/hyperactivity disorder. J Journal of the American Academy of Child Adolescent Psychiatry, 44(4), 377–384.

Masi, G., Millepiedi, S., Mucci, M., Bertini, N., Pfanner, C., & Arcangeli, F. (2006). Comorbidity of obsessive-compulsive disorder and attention-deficit/hyperactivity disorder in referred children and adolescents. Comprehensive Psychiatry, 47(1), 42–47.

Miller, G. A. (1956). The magical number seven, plus or minus two: Some limits on our capacity for processing information. J Psychological review, 63(2), 81.

Missonnier, P., Gold, G., Fazio-Costa, L., Michel, j.-p., Mulligan, R., Michon, A., et al. (2005). Early Event-Related Potential Changes During Working Memory Activation Predict Rapid Decline in Mild Cognitive Impairment. The Journals of Gerontology Series A: Biological Sciences and Medical Sciences, 60, 660–666.

Murata, T. (1989). Petri nets: Properties, analysis and applications. Proceedings of the IEEE, 77(4), 541–580.

Murdock Jr, B. B. (1962). The serial position effect of free recall. Journal of Experimental Psychology, 64(5), 482–488.

Nigg, J. T. (2006). What causes ADHD?: Understanding what goes wrong and why: Guilford Press.

Nouwens, S., Groen, M. A., & Verhoeven, L. (2017). How working memory relates to children’s reading comprehension: the importance of domain-specificity in storage and processing. J Reading writing, 30(1), 105–120.

O’Reilly, R. (2003). Making Working Memory Work: A Computational Model of Learning in Prefrontal Cortex and Basal Ganglia. Boulder: Institute of Cognitive Science.

O’Reilly, R. C., Braver, T. S., & Cohen, J. D. (1999). A Biologically Based Computational Model of Working Memory. In A. Miyake & P. Shah (Eds.), Models of Working Memory: Mechanisms of Active Maintenance and Executive Control (pp. 375–411). Cambridge: Cambridge University Press.

O’Reilly, R. C., & Frank, M. J. (2006). Making Working Memory Work: A Computational Model of Learning in the Prefrontal Cortex and Basal Ganglia. Neural computation, 18(2), 283–328.

Oberauer, K. (2009). Chapter 2 Design for a Working Memory. In Psychology of Learning and Motivation (pp. 45–100): Academic Press.

Oberauer, K., & Lewandowsky, S. (2011). Modeling working memory: a computational implementation of the Time-Based Resource-Sharing theory. J Psychonomic Bulletin Review, 18(1), 10–45.

Oberauer, K., Lewandowsky, S., Awh, E., Brown, G. D., Conway, A., Cowan, N., et al. (2018). Benchmarks for models of short-term and working memory. 144(9), 885.

Oberauer, K., Souza, A. S., Druey, M. D., & Gade, M. (2013). Analogous mechanisms of selection and updating in declarative and procedural working memory: Experiments and a computational model. Cognitive Psychology, 66(2), 157–211.

Page, M. P. A., & Norris, D. (1998). The Primacy Model: A New Model of Immediate Serial Recall. Psychological Review, 105(4), 761–781.

Peijnenborgh, J. C., Hurks, P. M., Aldenkamp, A. P., Vles, J. S., & Hendriksen, J. G. (2016). Efficacy of working memory training in children and adolescents with learning disabilities: A review study and meta-analysis. J Neuropsychological rehabilitation, 26(5-6), 645–672.

Peterson, L., & Peterson, M. J. (1959). Short-term retention of individual verbal items. Journal of Experimental Psychology, 58(3), 193–198.

Redgrave, P., & Gurney, K. (2006). The short-latency dopamine signal: a role in discovering novel actions? Nature Reviews Neuroscience, 7(12), 967–975.

Redick, T. S., & Engle, R. W. (2006). Working memory capacity and attention network test performance. J Applied Cognitive Psychology: The Official Journal of the Society for Applied Research in Memory Cognition, 20(5), 713–721.

Schweitzer, J. B., Lee, D. O., Hanford, R. B., Zink, C. F., Ely, T. D., Tagamets, M. A., et al. (2004). Effect of methylphenidate on executive functioning in adults with attention-deficit/hyperactivity disorder: normalization of behavior but not related brain activity. J Biological psychiatry, 56(8), 597–606.

Silva, M. (2013). Half a century after Carl Adam Petri’s Ph. D. thesis: A perspective on the field. J Annual Reviews in Control, 37(2), 191–219.

Sperling, G. (1960). The information available in brief visual presentations. Psychological monographs: General and applied, 74(11), 1.

Sperling, G. (1963). A Model for Visual Memory Tasks. Human factors, 5(1), 19–31.

Thiruvengada, H., & Rothrock, L. (2007). Affordance-based computational model of driver behavior on highway systems: A Colored Petri Net approach. In 2007 IEEE International Conference on Systems, Man and Cybernetics (pp. 888–893). IEEE.

Zhou, M., & Kurapati, V. (1999). Modeling, simulation, and control of flexible manufacturing systems: a Petri net approach. Singapore; [River Edge], NJ: World Scientific.

